# Breathing on Chip: Dynamic flow and stretch tune cellular composition and accelerate mucociliary maturation of airway epithelium *in vitro*

**DOI:** 10.1101/2021.05.07.443164

**Authors:** Janna C. Nawroth, Doris Roth, Annemarie van Schadewijk, Abilash Ravi, Tengku Ibrahim Maulana, Christiana N. Senger, Sander van Riet, Dennis K. Ninaber, Amy M. de Waal, Dorothea Kraft, Pieter S. Hiemstra, Amy L Ryan, Anne M. van der Does

## Abstract

Human lung function is intricately linked to blood flow and breathing cycles, but it remains unknown how these dynamic cues shape human airway epithelial biology. Here we report a state-of-the-art protocol for studying effects of physiological airflow and stretch on differentiation, cellular composition, and mucociliary clearance of human primary airway epithelial cells cultured on a perfused airway chip. Perfused epithelial tissue cultures developed a large airway-like cellular composition with accelerated maturation and polarization of mucociliary clearance when compared to traditional (static) culture methods. Additional application of airflow and stretch to the airway chip resulted in a cellular composition more comparable to the small(er) airways, reduced baseline secretion of interleukin-8 and other inflammatory proteins, and reduced gene expression of matrix metalloproteinase (MMP) 9, fibronectin, and other extracellular matrix factors. These results indicate that breathing-like mechanical stimuli are important modulators of airway epithelial cell differentiation and homeostasis and that their fine-tuned application could generate models of specific epithelial regions, pathologies, and mucociliary (dys)function.

## Introduction

The mucosal surface of the airways continuously filters contaminants from inhaled air and thereby provides an essential defense mechanism against infection and damage of the lungs. This function relies on an ensemble of specialized epithelial cells that use motile cilia to transport secreted mucus, along with trapped airborne matter, out of the airways^1^. Although static *in vitro* cultures of airway epithelial tissues feature all major cell types, full maturity of ciliary beat and mucociliary clearance is only reached after more than 60 days of differentiation^2^. Moreover, these culture conditions do not drive epithelial differentiation specifically towards the cellular composition characteristic of different airway branching generations, but rather depend on type of cell culture medium, donor, and culture protocol^3,4^. These limitations constrain scalability and translation of airway epithelial cell models, particularly for studying how infectious agents breach mature mucociliary clearance, or why some disease processes preferentially affect the large or the small airways^5^.

Given that mechanical stretch and shear stresses act on lung epithelial tissues throughout development and adult life *in vivo*^6^, differ in magnitude in large versus small airways^7,8^, and can improve airway epithelial barrier function *in vitro*^9^, we posited that differentiation, cellular composition, and ciliary function of human primary bronchial epithelial cell (hPBEC)-derived airway epithelial tissue may be modulated by physiologically relevant mechanical cues.

To test our hypothesis, we developed a microfluidic airway organ-chip protocol for differentiation of hPBEC at an air-liquid interface (ALI) on a flexible, porous membrane. Using this advanced cell culture system, we assessed differentiation and homeostasis of airway epithelium in response to a dynamic microfluidic environment with stretch and airflow shear stress levels that mimic the small airway environment during normal tidal breathing.

## Results

### Dynamic perfusion shifts airway epithelial cellular composition toward larger airways

To test our hypothesis, we used the microfluidic Chip-S1^®^ (Emulate Inc.), which features two adjacent, perfusable microchannels separated by a flexible and highly porous poly(dimethylsiloxane) (PDMS) membrane. hPBEC cultured on the apical side of the membrane can be exposed to air while interfacing fluidically with the (vascular) bottom compartment (Figure 1a top and Supplementary Figure 1a). The PDMS membrane is suited for dynamically stretching the epithelial tissue but was found to require extensive optimization of surface functionalization and cell culture protocol for successful hPBEC differentiation. Previously developed protocols for airway chips using cell-culture optimized, but rigid, membranes^10,11^ failed to reliably support cellular adhesion on the PDMS substrate (data not shown). After testing a variety of options, we found that coating the activated membrane with high concentrations of collagen IV (300 µg/mL), followed by seeding hPBEC at high density (∽300,000 cells/cm^2^), led to robust cell adhesion and confluent monolayer formation in the top channel. We subjected the hPBEC to dynamic medium perfusion in both top and bottom channel during the submerged phase and next switched to top air-liquid interface (ALI) with bottom channel medium perfusion to initiate differentiation (Figure 1a bottom). Consistent with prior studies^12^, the relatively large pore size of the membrane (7 µm) (Supplementary Figure 1b) stimulated spontaneous and progressive transmembrane migration of hPBEC in the majority of donors, as verified with our newly developed method of chip sectioning (Figure 1b left and Supplementary Figure c and d). Since bottom channel invasion limited visibility and frequently led to degeneration and delamination of tissue in the top channel, we evaluated a variety of strategies to prevent epithelial cell migration. Chemical cell migration inhibitors and changes to the cell culture medium (Supplementary Table 5) proved to be ineffective or detrimental to differentiation (data not shown), gel barriers were effective but unreliable (Supplementary Figures 1e and 2), and published endothelial barriers^13^ would require co-culture with additional cell types which was undesired for this study and in addition confounds the comparison to insert culture. We eventually identified a robust solution by functionalizing the bottom channel with the anti-fouling agent Pluronic F-127^14^. Using this biocompatible and removable anti-adhesion coating, bottom channel invasion was significantly reduced for all donors (Figure 1b right and Supplementary Figure 1f), enabling robust hPBEC differentiation as well as optional co-culture with primary endothelial cells at time point of choice (Supplementary Figure 3 and Supplementary Videos 1 and 2) as needed for applications involving vascular and immune cell responses^11^. Next, to assess the effects of the microfluidic and flexible environment on developing hPBEC tissues, we compared the perfused chip model to cell culture inserts, a standard static model, and evaluated hallmarks of airway epithelial differentiation, including pseudostratification and presence of epithelial cell types^3^ at day 14 of ALI^15^. Both chip and insert cultures exhibited a classical pseudostratified layer with basal cells located close to the membrane and luminal cells with visible ciliation (Figure 1c). Cell cultures in the chip exhibited significantly higher gene expression levels of basal cell markers (*TP63* and *KRT5*) and of keratin 8 (*KRT8*), a markers that is used for describing a population of cells for which there seems to be no consensus on its name yet as they are depicted in literature as non-basal^16^, luminal^16,17^, early progenitor^18^ or basal luminal precursor^19^ cells. For this reasons we will address them as KRT8^+^ cells. Expression of differentiated luminal cell markers for secretory club cell (*SCGB1A1*),goblet cell (*MUC5AC*) and ciliated cells (*FOXJ1*) was not different from insert cultures (Figure 1d). IF-staining of cell type-specific proteins confirmed similar levels of differentiated luminal cell populations with a (non-significant) increase in goblet cells in chip compared to insert cultures (Figure 1e and f).

**Figure 1.**
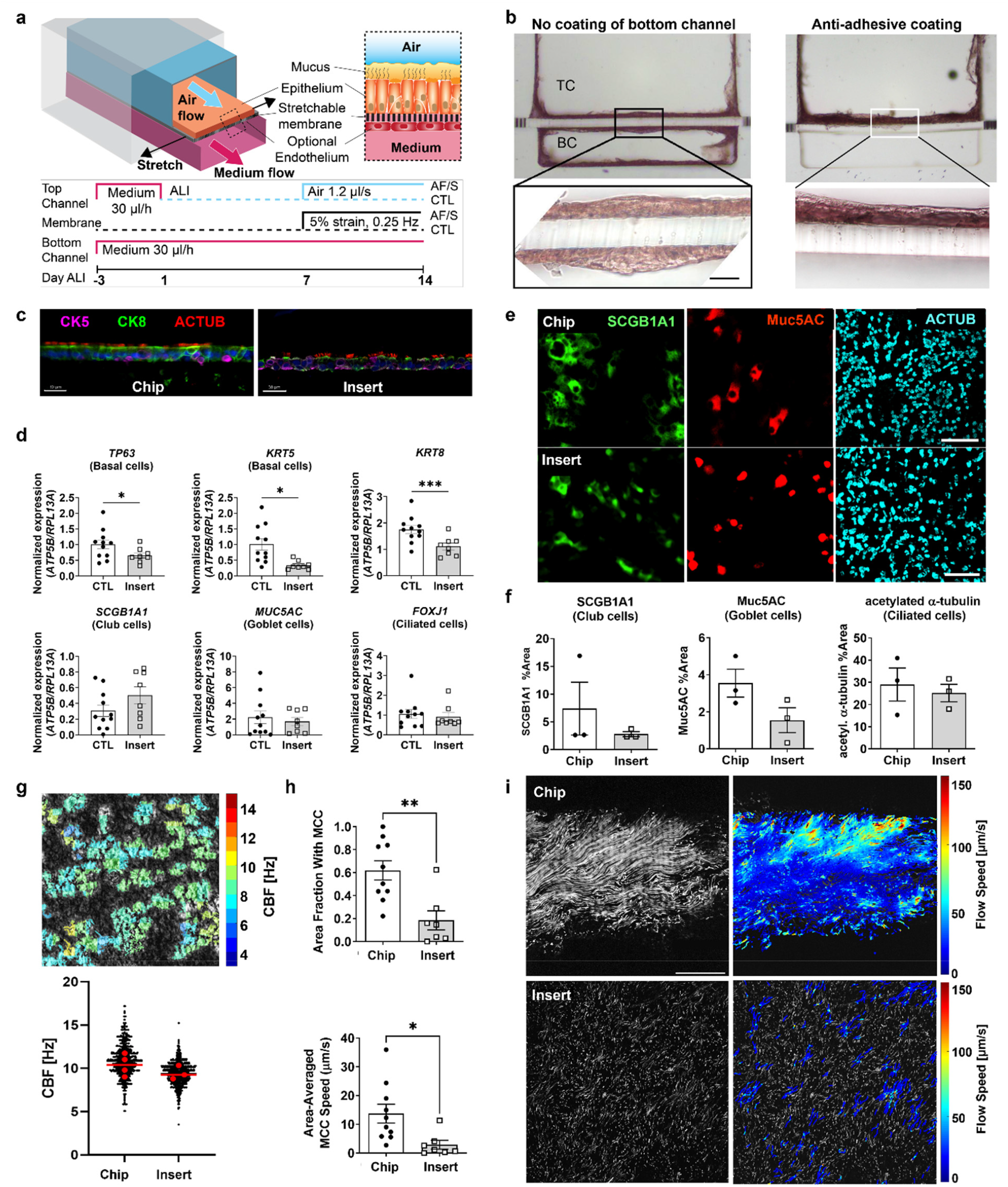
Dynamic chip culture skews cellular composition of the airway epithelium and accelerates maturation of mucociliary clearance in hPBECs. **a**, Airway basal epithelial cells are cultured in the top channel on a porous stretchable membrane. Endothelial cells can optionally be grown in the bottom vascular channel. Medium flow, airflow and membrane stretch are applied as indicated in the timeline. **b**, Histology cross-sections (H&E stain) of a fully invaded chip with hPBECs growing in both channels on the left, and a Pluronic treated chip with no invasion on the right. TC, top channel, BC, bottom channel. Zoom: cells line both sides of the membrane. Scalebar: 50 µm. **c**, Representative image of chip cross-section of a chip culture and an insert culture with stainings for cytokeratin 5 (CK5, basal cells), -8 (CK8), and acetylated alpha-tubulin (ACTUB, cilia). Scalebar: 30 µm. **d**, Gene expression of *TP63* (basal cells), *KRT5* (basal cells), *KRT8, SCGB1A1* (club cells), *MUC5AC* (goblet cells) and *FOXJ1* (ciliated cells) in collagen IV (COLIV)-coated chips and inserts at day 14 ALI. Data are shown as target gene expression normalized for the geometric mean expression of the reference genes *ATP5B* and *RPL13A*; N=8-11 donors, one chip or insert per donor. Data are depicted as mean +/-SEM. *, p<0.05; ***, p<0.001, a paired two-tailed paired t test. **e**, Representative IF staining of club (SCGB1A1), goblet (Muc5AC), and ciliated cells (acetylated alpha-tubulin, ACTUB) in chip and insert. Scalebar: 100 µm. **f**, Quantification of the cell culture surface area fraction positive for IF markers of each cell type. Depicted are mean +/-SEM from 3-6 biological samples from N=3 donors. No statistical difference was detected using one-tailed t test. **g**, Example of CBF heatmap (top; scalebar: 100 µm) and quantitative analysis (bottom). Data from N=2 donors, 2-4 chips or inserts per donor. Each black dot represents approximately 1 ciliated cell (chips: 2481 measurements; inserts: 2976 measurements). Red dots indicate means of individual inserts or chips and line indicates their median. **h**, Quantification of area fraction covered by MCC (left) and area-averaged MCC speed (right). Depicted are mean +/-SEM of one chip or insert per donor (chips N=9 donors; inserts: N=7 donors). *, p<0.05; **, p<0.01; two-tailed Welch’s t test. **i**, Example of fluorescent bead trajectories (left) and corresponding instantaneous flow speeds (right) in chip (top) and insert (bottom). Scale bar: 50 µm.

### Perfused airway epithelial tissue cultures show accelerated maturation of mucociliary clearance

Further exploring the differentiation status of the epithelium, we turned to functional markers of mucociliary clearance (MCC). High-speed videomicroscopy revealed a similar surface density and beat frequency of motile cilia in the two conditions (Figure 1g and Supplementary Video 3). We next directly assessed MCC function by tracking the displacement of fluorescent microbeads placed by the ciliated epithelium. Despite the similar beating frequency, we observed markedly better MCC in chip cultures compared to insert cultures exhibiting on average one third more MCC surface coverage and twice the tissue-averaged MCC speed (Figures 1h and I, Supplementary Video 4). It was previously shown that insert cultures lack efficient ciliary beat and MCC at day 14 of ALI, but markedly improve at around day 30 of ALI^2^. We measured MCC in cell culture inserts at day 35 and confirmed this improvement (Supplementary Figure 4, Supplementary Video 4), ruling out any underlying issues with the insert cultures and instead indicating that MCC maturation was accelerated on chip. Together, these observations suggest that culturing hPBEC on a flexible membrane combined with dynamic flow provides critical environmental cues that promote the differentiation and functional maturation of multiciliated airway epithelium *in vitro*.

### Dynamic airflow and cyclic strain shift airway epithelial cell composition toward smaller airways and polarize mucociliary clearance

Normal breathing creates a diverse mechanical landscape along the respiratory tree. Cyclic tissue strain tends to increase as a function of branching generation, whereas airflow shear stress reduces^7,8^. *In vitro* alveolar epithelial tissue responds to physiological cyclic strain by accelerated differentiation of specific cell types^20^. We hypothesized that breathing-related mechanical cues may also modulate differentiation of airway epithelia, potentially promoting the distinct epithelial phenotypes found in small versus large airways^3^. To test this hypothesis, we investigated the combined effect of small airway-like airflow in the top channel combined with cyclic membrane strain (AF/S) on hPBEC differentiation compared to control (CTL) chips in which no stretch was present and air exposure was static. The membrane of AF/S chips was actuated linearly and perpendicular to the channel at a rate of 0.25 Hz to achieve maximally 5% cyclic strain in the center of the chip, the estimated physiological strain in the small airways during tidal breathing at rest^7^. Airflow in AF/S chips was set to 1.2 µl/s, generating air flow shear stress of ca. 0.1 mPa on the cells, which is comparable to small airway conditions during tidal breathing^21^.

We applied AF/S between day 7 and 14 of ALI (Figure 1a bottom) to allow for barrier formation before the onset of mechanical stresses and provide sufficient time for key cellular markers to change in expression during the course of the experiment^15^. Like CTL chips, AF/S chips exhibited classical stratification with basal cells located near the membrane and differentiated luminal cells located near the surface (Figure 2a). We verified that lactate dehydrogenase (LDH) levels were similar between both chip groups, indicating comparable levels of cell viability (Supplementary Figure 6a). Compared to CTL chips, AF/S chips expressed significantly lower levels of genetic markers for basal cells (*TP63* and trend for *KRT5* (p=0.055)) and for KRT8^+^ cells. Expression of *SCGB1A1*, a marker of club cells, was significantly increased (Figure 2b). The difference in gene expression may capture an early stage in club cell development as the tissue-level density of immunofluorescent (IF) labeled club cell protein was similar between groups (Figure 2c and Supplementary Figure 6b). These results indicate that the AF/S stimulation may skew differentiation to a more small-airway like epithelial phenotype, which is characterized by a lower proportion of basal cells and higher proportion of club cells when compared to larger airways^3,4^.

**Figure 2.**
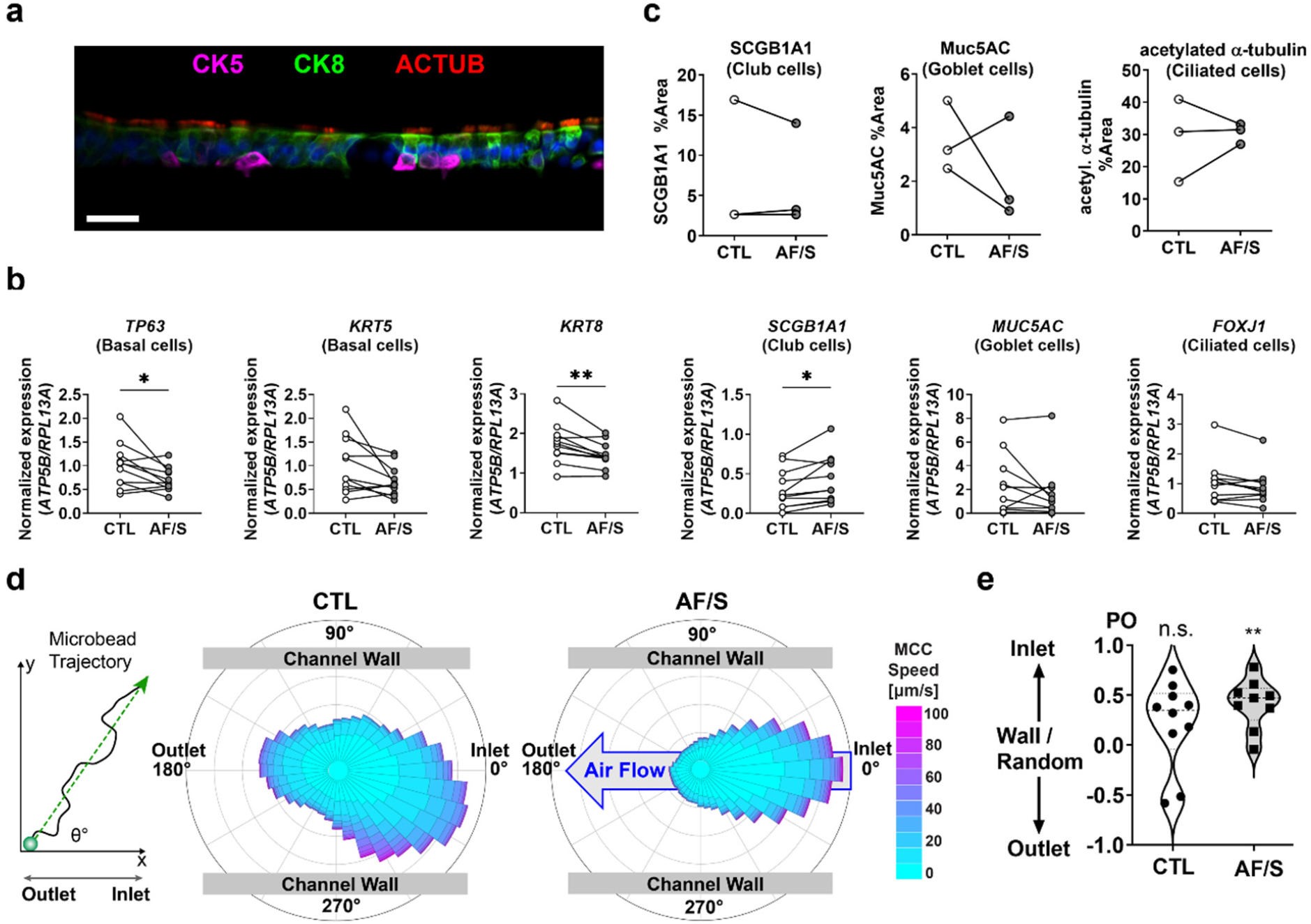
Exposure to airflow and stretch modulates cellular differentiation and polarizes mucociliary clearance in hPBEC. **a**, Representative image of chip cross-section of a chip culture exposed to airflow and stretch for 7 days fluorescently stained for cytokeratin 5 (CK5), -8 (CK8) or acetylated alpha-tubulin (ACTUB). Scalebar: 30 µm. **b**, Gene expression at day 14 ALI of *TP63* (basal cells), *KRT5* (basal cells), *KRT8* (differentiated non-basal cells), *SCGB1A1* (club cells), *MUC5AC* (goblet cells) and *FOXJ1* (ciliated cells). Open circles: CTL chips; gray circles: AF/S-exposed chips; N=8 donors, one chip per donor. Data are depicted as mean with SEM. *, p<0.05; **p<0.01 as assessed by a two-tailed paired t test. **c**, Quantification of the relative surface area labeled with IF markers for each cell type. N=3 donors; each point is mean value from 1 chip. N.S. as assessed by Wilcoxon matched-pairs signed rank test. **d**, From the overall transport direction of each fluorescent microbead (schematic on left), angle histograms of MCC direction for each condition are created, indicating the flow alignment relative to channel and applied flow (CTL: 140,760 trajectories; AF/S: 119,696 trajectories). Data from N=9 (CTL) and N=8 (AF/S) donors, 1 chip per donor. **e**, Polar order (PO) parameter obtained from the bead trajectories in (e). The PO ranges from -1 to 1, with -1 indicating perfect polarization towards the outlet and 1 towards the inlet, and 0 is random flow or towards the wall. Each data point is PO from 1 chip per donor with N=9 (CTL) and N=8 (AF/S) donors. **, p<0.01 as assessed by 1-sample Wilcoxon signed rank test against null hypothesis (random or wall-bound flow with PO=0).

Next, we assessed ciliary beat activity and mucociliary clearance function after AF/S application. We found no significant difference in the density of motile cilia, CBF, average MCC speed, or MCC coverage (Supplementary Figure 6c to e). We noted that MCC in all chips tended to be aligned along the length of the channel and point towards either end of the chip. However, whereas MCC in CTL chips only showed a weak bias towards the inlet, MCC in AF/S chips was most often directed towards the inlet, i.e., against the applied air and medium perfusion, consistent with the angle histograms of the fluorescent microbead trajectories for each condition (Figure 2d). We quantified the polarization of MCC by using a normalized index of direction, the polar order parameter (PO). The PO analysis confirmed unidirectional MCC towards the inlet in AF/S chips, but not in CTL chips, when compared to the null hypothesis of random or perpendicular flow direction (Figure 2e). By contrast, a recent study showed that human induced pluripotent stem cell (hiPSC) derived airway epithelium generated MCC in the same direction as imposed flow^22^. Hence, different mechanisms may be implicated in determining ciliary beat polarity of hPBEC versus hiPSC derived airway tissues. Together, our observations suggest that physiological levels of airflow and stretch may have profound effects on *in vitro* airway epithelial cellular composition and ciliary beat polarity.

### Dynamic airflow and cyclic strain affect developmental pathways and accelerate key steps of multiciliated cell maturation

To better understand the effects of dynamic culture on tissue organization and MCC function in hPBECs, we performed an unbiased bulk RNA-sequencing (RNAseq) analysis on RNA isolated from hPBEC in static cultures (inserts) and dynamic cultures (CTL and AF/S chips) from three different donors collected at day 14 of ALI. Principal component analysis (PCA) revealed a pronounced difference between the transcriptional profiles of inserts and chip cultures, and a smaller difference between donors (Figure 3a). We identified 182 differentially expressed genes (DEGs; q-value <0.05 and >1.5 fold change) between CTL chips and inserts (Supplementary Table 1) and performed pathway analysis (Top 10 depicted in Table 1, Supplementary Table 2, heatmaps in Figure 3b and Supplementary Figure 5). Interestingly, development-related gene expression was changed in chips when compared to cultures on inserts (Figure 3b), consistent with the observed larger proportion of differentiating cells and the more mature mucociliary clearance in chips compared to inserts. One of the pathways associated to the DEGs was tissue development. Compared to inserts, chip cultures also exhibited decreased expression of Wnt3a signaling-related (target) genes, including *WNT3A, AXIN2* and *FGF20*. Applying AF/S to the chip cultures shifted the transcriptional profile further (Figure 3c). We identified 20 DEGs between AF/S chips and CTL chips (Figure 3c, Supplementary Table 3). Further analysis did not reveal a specific pathway induced by airflow and stretch; however, several genes related to TGF-β/BMP signaling (i.e., *BMP2*; *INHBA* and *PDGFB*) were reduced in expression in AF/S compared to CTL (Figure 3c), which might play a role in the observed differences in cell type composition.

**Figure 3.**
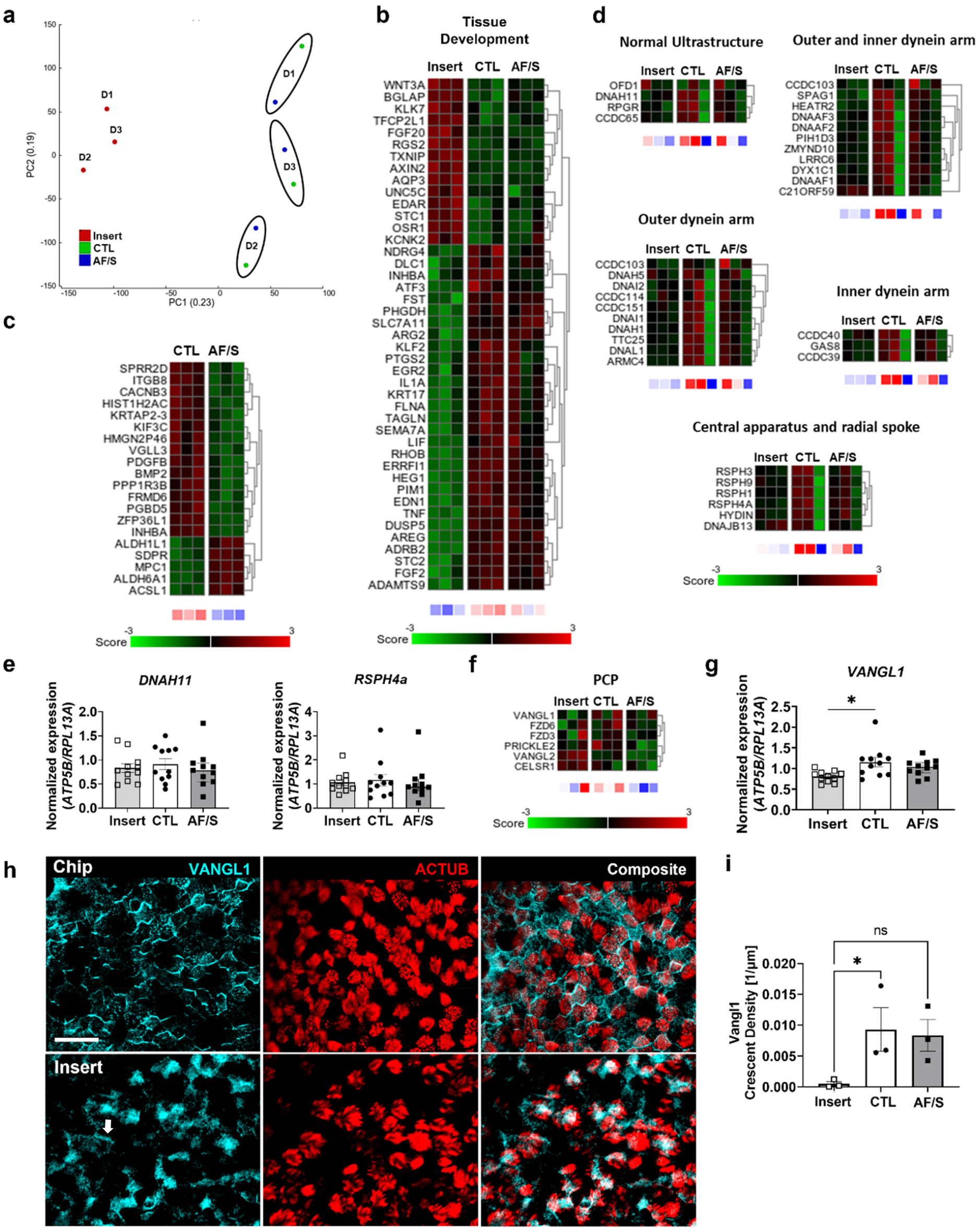
Culture on chip changes gene expression related to developmental pathways and promotes VANGL1 expression. **a**, PCA on all transcriptomes showing distinct clusters of airway epithelial cells from donors (N=3) in CTL chips, AF/S chips, or cell culture inserts coated with collagen IV. **b**, Heat maps displaying the Z score of DEGs related to the tissue development pathway depicted in Table 1, expression was compared between insert, CTL chip and AF/S chip cultures with N=3 donors that are paired (one donor per column). **c**, Heat map displaying the Z score of all 21 DEGs between AF/S and CTL chip cultures. N=3 donors that are paired between AF/S and CTL (one donor per column). **d**, Heat map displaying the Z score genes related to cilia structure and function between insert and chip CTL and AF/S cultures. **e**, Gene expression at day 14 ALI of *DNAH11* and *RSPH4a* for cultures on inserts (light gray bars; open squares), chips (open bars; black circles) or chips exposed to AF/S (dark gray; open circles); N=8-11 donors, one insert or chip per donor. Data are depicted as mean with SEM. **f**, Heat maps displaying the Z score of PCP genes between insert, CTL chip and AF/S chip cultures with N=3 donors that are paired between insert, CTL chip and AF/S chip cultures (one donor per column) and **g**, gene expression at day 14 ALI of *VANGL1* for cultures on inserts (light gray bars; open squares), chips (open bars; black circles) or chips exposed to AF/S (dark gray; open circles); N=8-11 donors, one insert or chip per donor. Data are depicted as mean with SEM. **h**, Representative IF staining of VANGL1 at day 14 ALI. Distinct VANGL1 crescents line most ciliated cells in chips of either condition (shown here: CTL). In contrast, static inserts of the same donor exhibit diffuse VANGL1 expression in most cells and very few such crescents (arrows). Scalebar 25 µm. **i**, Quantification of VANGL1 crescent density in CTL and AF/S chips and inserts at day 14 ALI. N=3 donors; each point is mean value from one chip per donor. *, p<0.05 as assessed by one-way paired ANOVA with Dunnett’s multiple comparison test.

**Table 1.** Top 10 Pathways associated with DEGs between CTL chip and insert cultures.

| Pathway description [# genes] | # DEGs | ID | p value | q value |
| --- | --- | --- | --- | --- |
| GOBP_RESPONSE_TO_OXYGEN_CONTAINING_COMPOUND [1653] | 43 | GO:1901700 | 2.76 e<sup>-22</sup> | 2.87 e<sup>-18</sup> |
| GOMF_SIGNALING_RECEPTOR_REGULATOR_ACTIVITY [543] | 27 | GO:0030545 | 4.15 e<sup>-21</sup> | 2.16 e<sup>-17</sup> |
| GOBP_TISSUE_DEVELOPMENT [1916] | 44 | GO:0009888 | 1.03 e<sup>-20</sup> | 3.56 e<sup>-17</sup> |
| GOBP_REGULATION_OF_CELL_POPULATION_PROLIFERATION [1745] | 41 | GO:0042127 | 1.23 e<sup>-19</sup> | 3.12 e<sup>-16</sup> |
| GOMF_MOLECULAR_FUNCTION_REGULATOR [1953] | 43 | GO:0098772 | 1.5 e<sup>-19</sup> | 3.12 e<sup>-16</sup> |
| GOBP_LOCOMOTION [1921] | 42 | GO:0040011 | 5.59 e<sup>-19</sup> | 9.7 e<sup>-16</sup> |
| GOMF_SIGNALING_RECEPTOR_BINDING [1550] | 38 | GO:0005102 | 8.29 e<sup>-19</sup> | 1.23 e<sup>-15</sup> |
| GOBP_REGULATION_OF_MULTICELLULAR_ORGANISMAL_DEVELOPMENT [1394] | 36 | GO:2000026 | 1.53 e<sup>-18</sup> | 1.99 e<sup>-15</sup> |
| GOBP_REGULATION_OF_CELL_D<br>EATH [1643] | 38 | GO:0010941 | 5.73 e<sup>-18</sup> | 6.62 e<sup>-15</sup> |
| GOBP_CELL_MIGRATION [1556] | 37 | GO:0016477 | 6.92 e<sup>-18</sup> | 7.19 e<sup>-15</sup> |

To probe the accelerated MCC maturation and polarization in dynamic chip culture conditions (Figure 1 & 2), we analyzed effects on genes related to cilia function, e.g., dyneins, in the RNAseq data. Compared to statics inserts, chip cultures from two out of three donors showed increased expression levels in this gene set (Figure 3d: organized based on literature^23^). Follow-up qPCR analysis of a selection of these genes using a larger number of donors revealed no significant changes in dynein-related genes *DNAH11* and *RSPH4a* (Figure 3e). We next assessed expression of planar cell polarity (PCP) genes (Figure 3f) and found that expression of PCP gene *VANGL1*, which plays an important role in multiciliated cell maturation^2,24^, was significantly increased in CTL chips compared to inserts (Figure 3g). We further investigated VANGL1 on protein level. IF staining at day 14 ALI showed that localization of VANGL1 in both chip conditions was asymmetric and appeared in a crescent shape along boundaries of ciliated cells, whereas expression was diffuse with few crescent shapes in insert cultures of same age (Figure 3h and i) but became comparable to chips at day 35 ALI (Supplementary Figure 7). Asymmetric PCP localization to cell boundaries is a key indicator of functional maturation of ciliary beat in airway epithelia^2,24^. Together, these results support that the enhanced MCC differentiation on chip might be promoted by accelerated maturation of multiciliated cells.

### Dynamic airflow and cyclic strain lower inflammatory IL-8 and reduce ECM-related gene expression in airway epithelial tissues

Studies focusing on mechanical ventilation have shown that *in vitro* application of short-term, non-physiological strain levels (e.g. up to 20% strain at 2 Hz for 30 minutes) triggers proinflammatory responses, such as *COX-2*/*PTGS2* mediated release of prostanoids^25^, and increased production of the inflammatory mediator IL-8 in lung epithelial cells^26–28^. In contrast, we found that stimulation with breathing-like AF/S at 5% strain at 0.25 Hz for 7 days led to downregulation of *PTGS2* by as shown by RNAseq (Figure 3b) and qPCR analysis (Figure 4a). Furthermore, over the course of 7 days, IL-8 protein release in AF/S chip culture medium significantly decreased compared to CTL chips (Figure 4b). Since CTL and AF/S chips both exhibited normal morphologies, comparable ciliary function, and similarly low LDH levels (Supplementary Figure 6a), we concluded that the reduced IL-8 levels in AF/S chip cultures were not due to lower cell viability. To assess if baseline levels of additional cyto-/chemokines were affected, we used a multiplex assay. Among all tested proteins, IL-9 and MIP-1β were significantly reduced, and IL-8 and G-CSF were near significantly reduced (Figure 4c).

**Figure 4.**
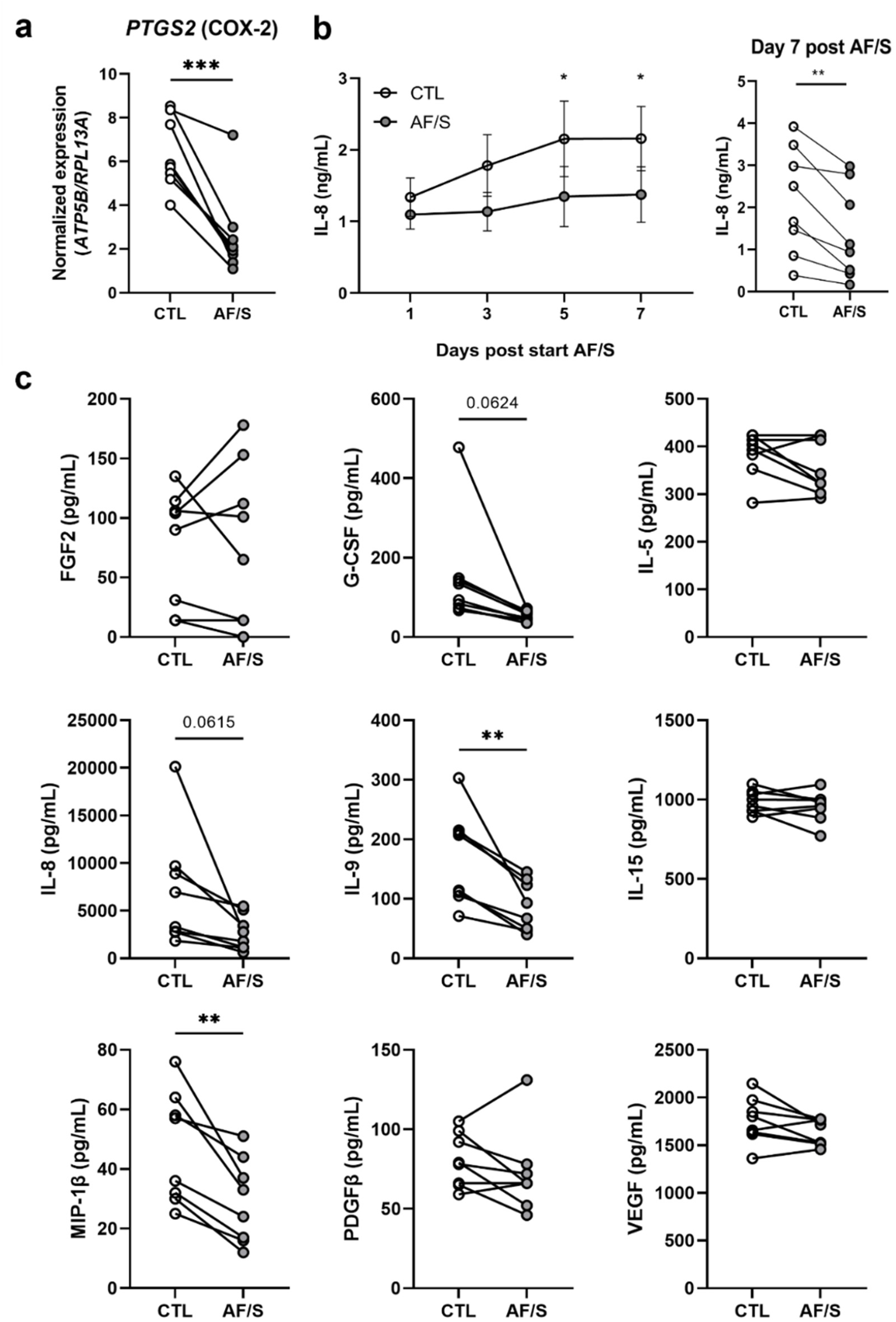
Exposure to airflow and stretch reduces pro-inflammatory baseline gene and protein levels. **a**, Gene expression of *PTGS2* at day 14 ALI. Open circles: CTL chips, gray circles: AF/S-exposed chips; N=8 different donors, one chip/donor. Data are depicted as mean +/-SEM. **, p<0.01 as assessed by paired t test. **b**, Left: IL-8 protein levels (ELISA) in the basal channel medium collected 24 h after each medium change at the indicated times. Filled circles are CTL chips, open circles are AF/S chips; N=8 donors with 2 chips per donor, except for 2 donors with 1 AF/S chip each. Right: paired IL-8 levels at day 14 post ALI and 7 days of application of AF/S. *, p<0.05 as assessed by Mixed-effects analysis combined with a Sidak’s multiple comparisons test. **, p<0.01 as assessed by paired t test. **c**, Baseline protein levels of FGF2; G-CSF; IL-5; IL-8; IL-9; IL-15; MIP-1β; PDGFβ and VEGF as assessed by multi-protein array analysis. N=8 donors, one chip per donor. **, p<0.01 as assessed by paired t test.

### Dynamic airflow and cyclic strain reduce production of extracellular matrix-related genes in airway epithelial tissues

Excessive mechanical stresses associated with mechanical ventilation have also been associated with extracellular matrix (ECM) remodeling through the upregulation and increased deposition of matrix components like collagens and fibronectin as well as their breakdown, for example by MMPs^29^. Our RNAseq data indicated that expression of specific ECM related genes was also altered in AF/S chips, including collagens, glycoproteins (selection), serpins, MMPs, and Lysyl oxidase homologs (LOXLs)-related genes (Figure 5a). Contrasting the effects seen in mechanical ventilation models^29,30^, TGF-β signaling, an activator of matrix remodeling, and expression of *MMP1, MMP9, COL1A1* (collagen type-1) and *FN1* (Fibronectin), were downregulated in AF/S compared to CTL (Figure 3c and Figure 5b). Except for TGF-β signaling, the effects were not significant for the low donor number tested in the RNAseq (N=3); however, qPCR analysis of *MMP9* and *FN1* for N=8 donors demonstrated significant downregulation of both genes in AF/S chips compared to CTL chips (Figure 5c). Together, these results suggest that physiologically relevant mechanical cues may downregulate proinflammatory signaling and ECM-remodeling in airway epithelial tissues.

**Figure 5.**
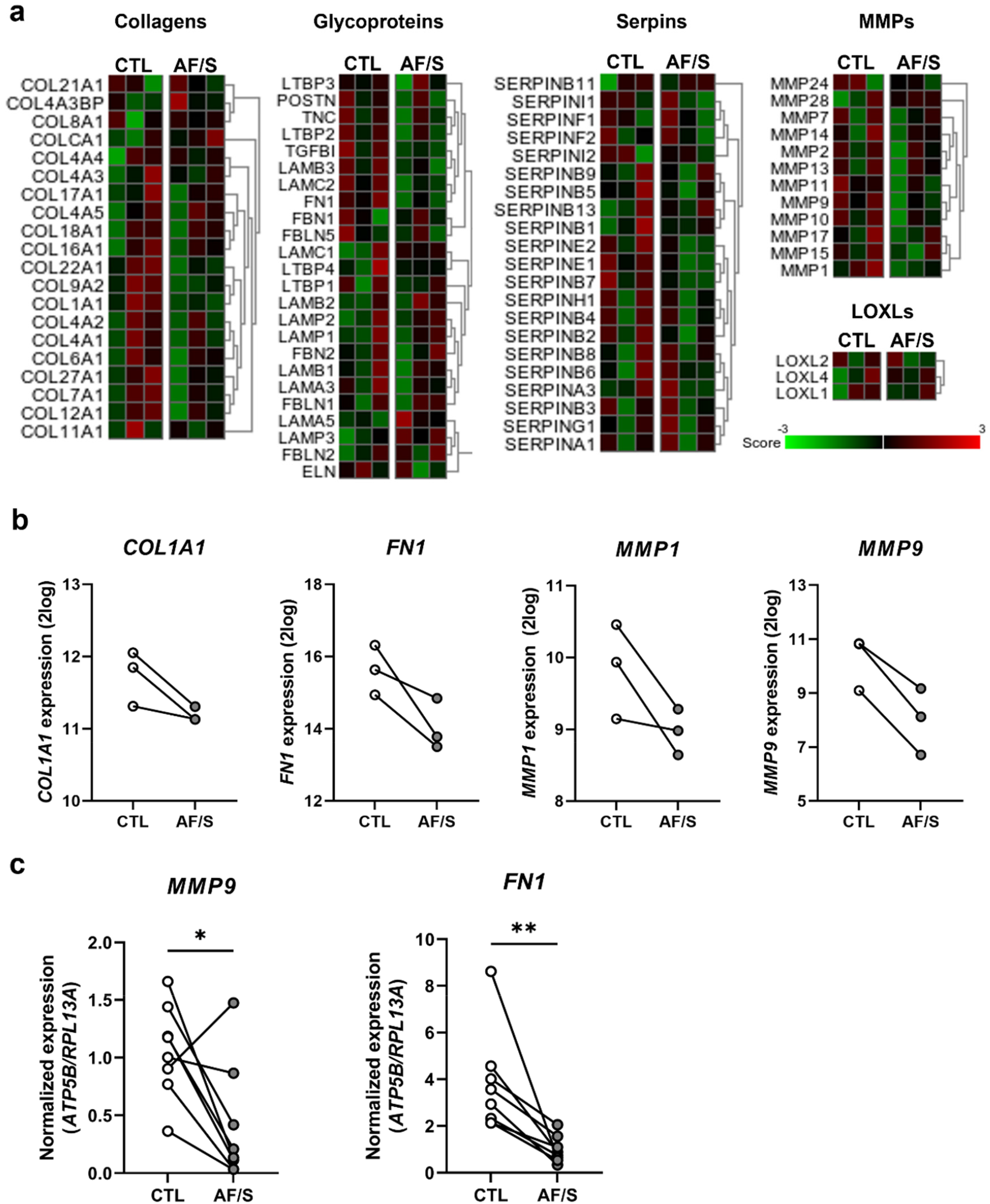
Exposure to airflow and stretch is associated with downregulation of ECM-related genes. **a**, Heat map of Z-scores for ECM-related gene expression (RNAseq), i.e., collagens, glycoproteins, matrix metalloproteinases (MMPs), SERPINs and LOXLs, in AF/S and CTL chips at day 14 of ALI. RNA was derived from N=3 donors that are paired between AF/S and CTL (one column per donor)), 1 chip/donor. **b**, Gene expression derived from the RNAseq dataset of selected genes shown in (A), i.e., collagen type-1 (*COL1A1*), fibronectin (*FN1*), matrix metalloprotease 1 and 9 (*MMP1* and *MMP9*) by AF/S and CTL chip cultures at day 14 of ALI. **c**, Gene expression assessed by qPCR analysis of *MMP9* and *FN1* at day 14 of ALI, Open circles: CTL chips, gray circles: AF/S chips.; *p<0.05; **, p<0.01 as assessed by a paired t test, N=8 donors, one chip per donor.

## Discussion

Primary human airway epithelial cells differentiated in perfused chips exhibited a notable increase in a development-related gene signature, showed enhanced mucociliary clearance, and developed a cellular composition more characteristic of the larger airways (e.g. greater proportions of basal and goblet cells^31^) compared to static culture. Chip cultures exhibited altered expression of genes related to variable pathways, including development. The unchanged levels of the other differentiated luminal cells and accelerated maturation of mucus clearance suggest a physiological role of the development-related pathways on chip that may contribute to a large airway-like phenotype. Additional exposure to small airway-like levels of air flow shear stress and cyclic stretch (AF/S) reduced expression of these pathways as well as inflammation and ECM-remodeling related genes and led to a transition towards an epithelial cell-type composition resembling the small(er) airways (e.g. lower basal and higher club cell levels^31^). Interestingly, the AF/S stimulation polarized MCC towards the applied flow direction, indicating an impact on tissue-wide multiciliated cell organization^32^. Together, these results suggest that stresses from physiological levels of flow and stretch directly modulate the phenotype and mucociliary maturation of airway epithelia; however, other mechanochemical cues that differ between chip and static culture, such as cellular substrate, mucus rheology, or medium mixing may also contribute. Regarding potential mechanisms, we hypothesize that flow and stretch modulate the observed on-chip activation of developmental pathways, such as EGF, BMP, FGF and Wnt signaling, which are known to influence airway epithelial cell fates. For example, EGF/EGFR signaling is involved in the distal to proximal repatterning of the smaller airways in smokers^5^. Our hypothesis is supported by other studies that propose a role of mechanical cues in determining airway epithelial cell fate: YAP/TAZ signaling, well-known to be triggered by mechanical cues, inhibits goblet cell differentiation in mice^33^. YAP or TAZ gene expression was not changed in our RNAseq dataset; however, any acute response may have faded after the 7 days of exposure to airflow and stretch. The observed changes in for example inflammatory gene expression could also result from changed cellular composition, for which single cell RNAseq analysis would be an interesting tool to further unravel these effects. Taken together, we propose that the proper tuning of developmental pathways by mechanical stimuli can skew differentiation to either proximal or distal patterning. The methods we developed will support the development of better modeling of lung tissue compartments *in vitro* and enable future experiments into the mechanisms by which this pathway activation, and resulting patterning goes awry in disease or damage, and whether such maladaptive airway remodeling can be reversed by application of targeted mechanical cues.

In this study we used human primary cells from multiple human donors (Supplementary Table 4) and studied differentiation in the presence of mechanical stimulation in order to capture robust, longer-term responses rather than acute insults or cell-line specific behaviors. We did not co-culture endothelial cells in the vascular compartment of the chip in this study as this would confound the effect of biomechanical cues on airway epithelial biology. It would be interesting to study in the future how the changes in epithelial cellular composition affect the mesenchymal and the vascular compartment. Although we did not explore the isolated effects of dynamic flow and stretch on airway epithelial biology, the ability to deliver these cues and achieve measurable effects on airway differentiation and homeostasis using a commercially available chip model combined with a robust protocol for primary airway culture opens new avenues for investigating ciliogenesis, airway disease and epithelial regeneration in the presence of mechanical stresses and establish beneficial versus adverse effects. Our method could also be applicable to the differentiation of alveolar or airway cells derived from induced pluripotent stems cells^22,34^ as well as multi-tissue airway models and enhance their structural and functional maturation *in vitro*.

## Acknowledgements

We would like to acknowledge Sowmya Balasubramanian, Marianne Koliana, Ben Calvert, and Erik Quiroz for their help with cell culture and chip maintenance. We thank the Emulate team, including Geraldine Hamilton, Carolina Lucchesi, Antonio Varone and Lorna Ewart, for support with Emulate equipment and technical discussions. We thank Lennart Voortman, Annelies Boonzaier-van der Laan and Mees de Graaf for testing imaging of the chip and training on the Dragonfly microscope and Tom Ottenhoff and Simone Joosten for the Luminex assay. The work at LUMC is supported by an EU Marie Curie Global Fellowship (#748569), the Dutch Society for the Replacement of Animal Testing (Stichting Proefdiervrij), the Netherlands Organization for Health Research and Development (ZonMw; #114021508) and the Lung Foundation Netherlands (grant #4.1.19.021). ALR is supported by the Hastings Foundation, the Tyler Health and Education Trust and Cystic Fibrosis Foundation (grants #FIRTH17XX0 and #FIRTH21XX0). JCN is supported by an ERC-STG (MecCOPD).

## Methods

### Primary Human Bronchial Epithelial Cell Sourcing and Expansion

Here and in following steps, we report the procedures optimized in parallel in laboratories at LUMC, USC, and Emulate Inc. and specify differences in material and protocol choices as applicable. For donor characteristics see Supplementary Table 4.

#### Leiden

hPBEC were isolated from macroscopically normal lung tissue obtained from patients undergoing resection surgery for lung cancer at the Leiden University Medical Center, the Netherlands. Patients from which this lung tissue was derived were enrolled in the biobank via a no-objection system for coded anonymous further use of such tissue (www.federa.org). However, since 29-11-2020, patients are enrolled in the biobank using active informed consent in accordance with local regulations from the LUMC biobank with approval by the institutional medical ethical committee (B20.042/Ab/ab and B20.042/Kb/kb). hPBEC were thawed in a T75 flask coated with 30 μg/mL bovine type-1 collagen (PureCol^®^, Advanced BioMatrix), 10 μg/mL fibronectin (Promocell, PromoKine, Bio-connect) and 10 μg/mL BSA (ThermoFisher Scientific) in BEpiCM-b basal medium (ScienCell Research Laboratories, Sanbio), supplemented with 100 U/mL penicillin (Lonza), 100 μg/mL streptomycin (Lonza) and bronchial epithelial cell growth supplement (ScienCell). When reaching ∽90% confluency, cells were trypsinized in 0.03% (w/v) trypsin (ThermoFisher Scientific), 0.01% (w/v) EDTA (BDH, Poole, UK), 0.1% glucose (BDH) in phosphate buffered saline (PBS) and prepared for seeding.

#### USC

Fresh hPBEC were isolated from lung explant tissue from rejected donor transplant from subjects with no prior evidence of chronic lung disease with IRB approval from the University of Southern California (IRB# USC HS-18-00162) using established protocols^35^. Next, hPBEC were thawed in T75 flasks or 100 mm cell culture dishes coated with PureCol (Advanced Biomatrix, Carlsbad, CA, USA) using complete BEpiCM-b basal medium as described above. When reaching ∽90% confluency, cells were dissociated with Accutase (Sigma Aldrich, St. Louis, MO, USA) and prepared for seeding.

#### Emulate Inc

Hydrogel experiments were performed with hPBECs from a commercial supplier (Lifeline Cell Technologies, Frederick, MD, USA) that were cultured similar to the methods stated for LUMC/USC.

### Chip functionalization and cell culture

#### Chip membrane activation

To hydrophilize the PDMS membrane, top and bottom channels of the Chip-S1^®^ (Emulate, Inc., Boston, MA, USA) were filled with ER-1 solution (1 mg/mL in ER-2; Emulate Inc) followed by 10 minutes of UV-light exposure using a 36-Watt UV-chamber (NailStar Professional, Model NS-01-US). Next, channels were washed twice with ER-2 (Emulate Inc.). This ER-1/ER-2 procedure was repeated once more. After the second ER-2 wash, channels were washed and filled with cold PBS.

#### Chip membrane coating

The chip top channel was emptied and subsequently filled with 300 µg/mL human collagen IV (Sigma) solution in PBS. Chips were placed in petri dishes containing a small, open container of PBS to promote humidity and kept overnight in the incubator at 37°C/5%CO_2_. For optimization experiments, also other coating components and their mixtures were used: 0.1% BSA (Gibco), 30 µg/mL bovine type-1 collagen (PureCol^®^, Advanced BioMatrix), or human fibronectin (Sigma). We found that BSA and collagen IV coating both resulted in reliable cell adhesion; however, collagen IV coated chips exhibited significantly better viability post seeding and significantly higher ciliation compared to BSA coated chips at day 14 ALI (Supplementary Figure 8).

#### Epithelial cell seeding and submerged culture on chip

The coating solution in the top channel was removed and replaced with so-called B/D complete medium: a 1:1 mixture of BEpiCM-b and DMEM medium (STEMCELL Technologies), supplemented with Bronchial Epithelial Cell Growth Supplement (ScienCell), and additional 50 nM EC-23 (Tocris); 25 mM HEPES (Cayman Chemical), 100 U/mL penicillin and 100 µg/mL streptomycin (ScienCell). B/D complete was pre-filtered using a 70 µm cell strainer (BD Biosciences) or a 0.22 µm vacuum filter unit (Steriflip™; Millipore, Sigma) to remove any suspended particles that could block flow in the Chip-S1^®^. Epithelial cells were seeded in B/D complete in the top channel at 3×10^6^ cells/mL (ca. 90K cells per chip, i.e., ca. 320K cells/cm^2^) and left to adhere in the incubator for ∽6h. At this point, the Pluronic treatment described below was applied to the bottom channel in the final chip protocol. Next, chip channels were washed gently with pre-warmed B/D complete. The chips were connected to the pre-warmed media-filled fluidic manifolds, “Pod^®^” (Emulate Inc.). After obtaining fluid connection between chips and Pods, these units were placed in the micro perfusion instrument, “Zoë^®^” (Emulate Inc.). After finishing the initial regulate cycle program (which pressurizes the medium to increase gas solubility and remove nucleating air bubbles while the system calibrates), the chips were continuously perfused with a flow rate of 30 µl/h in both top and bottom channel. Approximately 24h after start of the first regulate cycle, a so-called via wash was performed, dislodging any bubbles in the Pod’s reservoirs fluid vials, followed by a second regulate cycle. This was important to minimize the blockage of flow by air bubbles. When following this protocol, no degassing of medium was needed. Approximately 3-5 days after seeding (donor dependent), air-liquid interface (ALI) was established as described in the next paragraph.

#### Epithelial differentiation at air-liquid interface (ALI) in the airway chip

ALI was established by removing the medium from the top channel and having only medium flowing through the bottom channel. To equalize (hydrostatic) pressure at the membrane level, reservoirs in the Pod connecting the inlet and outlet of the top channel were sealed with 1 mL B/D complete medium per reservoir. Nonetheless, occasional flooding of the top channel would still occur, requiring daily monitoring. Submerged top channels were emptied manually using a 1000 µl pipette. Epithelial cultures were also monitored daily for flow issues and cellular invasion and morphology. If flow issues arose, chips were disconnected, channels were rinsed with medium, potential air bubbles were dislodged, and chips were re-reconnected after establishing liquid-liquid interface between chip and Pod. If this happened repeatedly in the same sample, the Pod was replaced. Every 48h, medium outlet reservoirs were emptied, and inlet reservoirs were filled with fresh medium.

#### Application of airflow and stretch

At day 7 of ALI, the nominal flow rate of the top channel in AF/S chips was set to 120 µl of medium per hour which corresponds to ca. 1.2µ/s of airflow. Additionally, the membrane strain rate was set to 5% at a frequency of 0.25Hz.

### Invasion scoring

We assigned the following scores to the degree of cell invasion into the bottom channel, as assessed by phase contrast microscopy: 0, no invasion; 1, individual cells in the bottom channel; 2, confluent cell layers in the bottom channel; 3, confluent tissue with multiple layers and/or fibroblast-like cell morphology in the bottom channel.

### Prevention of bottom channel invasion

#### Chemical treatments

Several strategies were tested to assess inhibition of PBEC migration to the bottom channel which are summarized in Supplementary Table 5. Three strategies were included: cell culture media variations, addition of different dose of EC-23 to the currently used medium or addition of dexamethasone at variable concentrations to the cell culture medium. Alternatively, several inhibitors or agonists of signaling pathways were added to the cell culture medium currently used. Neither signaling inhibitors of migration nor altered media composition significantly reduced invasion while preserving differentiation capacity (data not shown).

#### Hydrogel mechanical barrier

The concurrent seeding of epithelial and endothelial cells onto either side of the membrane can be used as a mechanical barrier to cell migration^36^, however, this strategy prevents the study of epithelial cells in isolation and comparison to standard culture conditions. Instead, we tested the use of a mechanical gel barrier. Different compositions of ECM gels in the top channel were tested, including bovine type-1 collagen (FibriCol^®^, Advanced BioMatrix), Matrigel^®^ (Corning, Corning, NY, USA), Fibronectin (Gibco), collagen IV (Sigma), and the collagen cross-linking agent, microbial transglutaminase (MTG) (Modernist Pantry LLC, Eliot, ME, USA). All ECM mixtures were prepared in PBS-solution and contained the following: collagen I = 0.5 mg/mL collagen I; collagen I + MTG = 0.5 mg/mL collagen I + 4 mg/mL MTG; collagen I + Fibronectin = 0.5 mg/mL collagen I + 200 µg/mL Fibronectin; collagen I + collagen IV = 0.5 mg/mL collagen I + 200 µg/mL collagen IV; collagen IV + Matrigel^®^ = 400 µg/mL collagen IV + 200 µg/mL Matrigel^®^. The gel was prepared by injecting the gel prepolymer solution after membrane surface activation and incubated overnight at 37°C. The next day, the gel was then flushed twice with 100 µL of warm medium at 187.5 µL/s flow (Eppendorf Xplorer automatic pipet; Eppendorf). This generated a 3D-ECM layer on the membrane with a thickness of 20-85 µm, as optimized previously^37^ (Supplementary Figure 2a and b). To assess and confirm ECM scaffold formation of different ECM compositions, we stained the gels with a fluorescent dye. Briefly, 1 mg/mL of N-hydroxysuccinimide (NHS) ester dye (Atto 488 NHS Ester, Sigma) was mixed with 50 mM borate buffer (pH 9) in 1:500 ratio. Directly after ECM formation, the prepared staining solution was injected into the top channel and the chips were incubated for 25 minutes at room temperature in the dark. The top channel was then rinsed three times with PBS prior to fluorescence imaging. Using ImageJ^38^, the ECM-covered area was measured as a percentage to the total channel area via automated thresholding. The entire straight length of the channel was evaluated in all conditions (N=3 chips each). To measure the ECM thickness, z-stack images with 4 µm step size were acquired in two defined regions (left and right part) of the top channel. Using ImageJ, the stacks were 3D-projected and rotated to obtain the 90°-side view of the channel, followed by visualizing the stained ECM with different thicknesses via automated thresholding. Average ECM thickness was measured by calculating the threshold area divided by the length of the image. Basal cells seeded onto the different gel types attached well, and significant reduction in basal channel invasion was observed compared to chips without hydrogels (Supplementary Figure 1e). At day 14 of ALI a well-differentiated epithelium containing club, ciliated, goblet and basal cells had formed (Supplementary Figure 2c). However, given substantial variability between ECM lots used for hydrogel preparation and other factors influencing spreading, gelling, and final geometry in the microfluidic channel, it was technically challenging to reliably deposit gels without any holes that allowed for transmigration.

#### Pluronic anti-adhesion surface treatment

Approximately 6 h after hPBEC seeding into the top channel, the bottom channels of all chips were coated with the surfactant Pluronic F-127 (Sigma) to prevent cell adhesion. For this, a minimum of 0.02% w/v Pluronic F-127 was freshly prepared in complete B/D medium and given time to dissolve at room temperature rather than at 37°C, as solubility of Pluronic decreases with temperature^39^. Next, the solution was sterile filtered using a 0.22 µm filter. The bottom channel of the chips was emptied and filled with the Pluronic solution, followed by 1h incubation at 37°C. After the incubation, warm medium was used to gently rinse both channels before proceeding with connecting the chips to the perfusion instrument Zoë as described above. Our preliminary data suggest that the bottom channel functionalization can be safely renewed at later timepoints by adding up to 0.2% Pluronic to the perfusion medium for 24 to 48h, which prevents bottom channel invasion even in highly migratory donors.

### Endothelial-epithelial co-culture on chip

Primary Human Lung Microvascular Endothelial Cells (HLMVEC) were purchased from Cell Applications (San Diego, CA, USA) and expanded in Microvascular Endothelial Cell Medium (Cell Applications). Chips were seeded with endothelial cells on day 11 or 12 of ALI. To prepare for endothelial cell seeding, the chips with epithelial tissue in the top channel were disconnected from the Pods. Since Pluronic is more soluble in medium at lower temperatures^39^, we washed the basal chip channels at day 12 of ALI with ice-cold buffer. For this, the top channel was filled with warm B/D complete medium, and the bottom channel was incubated three times for 30 seconds, then rinsed, with ice-cold PBS to remove the Pluronic coating. The bottom channel was filled with 100 µg/mL human fibronectin (Sigma) in B/D complete medium. The chips were flipped upside down, leveled and incubated for 1 h at 37° C. Just before endothelial cell seeding, cells were washed with PBS and dissociated using Accutase (Sigma), then resuspended at 7.5 ×10^6^ cells/mL in endothelial cell medium. The fibronectin solution was removed from the bottom channel and ca. 8 µl of the cell suspension was added, resulting in a seeding density of ca. 200, 000 cells/cm^2^. The chip was again flipped upside down and left undisturbed in the incubator for 2h. After 2h, the endothelial cell medium was carefully exchanged in the bottom channel. The chip was reconnected to the Pod and brought to ALI using standard procedures. The bottom channel was perfused with a mix of 50% endothelial cell medium and 50% B/D complete medium at standard flow rates (30 µl/h).

### Airway cell culture inserts

#### Leiden

To match the seeding conditions on the chips, hPBEC were seeded at high density (150,000 cells/well) on ECM-coated 12 mm clear polyester cell culture inserts with 0.4 μm pore size (Corning Costar, Cambridge, USA). Inserts were coated with 300 µg/mL human collagen IV (Sigma) solution in PBS. We also tested a coating mixture consisting of 30 μg/mL Purecol (Advanced BioMatrix,), 10 μg/mL fibronectin (Promocell) and 10 μg/mL BSA (ThermoFisher Scientific) in PBS^40,41^ but no significant differences in any our endpoints compared to the collagen IV-coated inserts (Supplementary Figure 10), hence we discontinued this alternative protocol.

Seeded cells were cultured submerged in B/D complete with EC-23 (50 nM). When cells were confluent, medium on the top side was removed to initiate differentiation at air-liquid interface. Medium was changed 3 times a week and the top side was washed with warm PBS.

#### USC

Clear polyester 6.5 mm cell culture inserts with 0.4 μm pore size (Corning Costar, Cambridge, USA;) were pre-coated with 300 µg/mL human collagen IV (Sigma) solution in PBS. hPBEC were seeded at high density (70.000 cells/well) and cultured as above.

### RNA isolation and bulk RNA sequencing

At day 14 of ALI, cells in the airway chip were lysed by disconnecting the chip from the Pod, lodging an empty filter tip at the outlet of the chip, and pipetting 100 µl lysis buffer (Promega) with another filter tip into the inlet of the chip. This tip was then lodged into the inlet, such that the two tips inserted at either channel end were capturing the overflowing liquid. When visual inspection indicated lysis of the cells, the lysis solution was collected in a tube. Another 100 µl was used to rinse the channel once more, to obtain in total 200 µl of lysis solution from one chip. The lysis solution was stored at -20°C until RNA extraction. Total RNA was robotically extracted using the Maxwell tissue RNA extraction kit (Promega) and quantified using the Nanodrop ND-1000 UV-Vis Spectrophotometer (Nanodrop technologies, Wilmington, DE, USA). Next, RNA was sent to GenomeScan (Leiden, the Netherlands) for Bulk RNA sequencing, or converted to cDNA for qPCR analysis. RNA sequencing was performed with the cDNA fragment libraries by the Illumina NovaSeq6000 sequencer using 150 bp pair-end sequencing settings.

### RNAseq analysis

#### Data processing

Data processing was performed by GenomeScan. For this, the NEBNext Ultra II Directional RNA Library Prep Kit for Illumina was used to process the samples. The sample preparation was performed according to the protocol “NEBNext Ultra II Directional RNA Library Prep Kit for Illumina” (NEB #E7760S/L). Briefly, mRNA was isolated from total RNA using the oligo-dT magnetic beads. After fragmentation of the mRNA, a cDNA synthesis was performed. This was used for ligation with the sequencing adapters and PCR amplification of the resulting product. The quality and yield after sample preparation were measured with the Fragment Analyzer. The size of the resulting products was consistent with the expected size distribution (a broad peak between 300-500 bp). Clustering and DNA sequencing using the NovaSeq6000 was performed according to manufacturer’s protocols. A concentration of 1.1 nM of DNA was used. Sequence reads were trimmed to remove possible adapter sequences using cutadapt v1.10. Presumed adapter sequences were removed from the read when the bases matched a sequence in the adapter sequence set (TruSeq adapters). For each sample, the trimmed reads were mapped to the human GRCh37.75 reference sequence (Homo_sapiens.GRCh37.75.dna.primary_assembly.fa). The reads were mapped to the reference sequence using a short-read aligner based on Burrows-Wheeler Transform (Tophat v2.0.14) with default settings. Based on the mapped locations in the alignment file the frequency of how often a read was mapped on a transcript was determined with HTSeq v0.11.0. The hit counts were summarized and reported using the gene_id feature in the annotation file. Only unique reads that fall within exon regions were counted. To enable comparison of gene/transcript expression across all samples outside of the context of differential expression analysis, RPKM/FPKM (reads/fragments per kilobase of exon per million reads mapped), TPM (transcripts per million), and CPM values were calculated with Cufflinks v2.2.1.

#### Differential expressed genes

The read counts were used in the DESeq2 v2-1.14 package, a statistical package within the R platform R v3.3.0. The differentially expressed genes were identified based on read counts of genes >10, with a log2 fold change of >1.5 and q value of <0.05.. Since groups had a small sample size and over-dispersion, an expression curve model based on a negative binomial distribution and local regression was used to estimate the relationship between the mean and variance of each gene. For heat maps, log2 expression of read counts are normalized with Z scores calculated by: (X (value of the individual sample) - μ (average of the row)) / s (standard deviation of the row).

#### Gene set analysis

The gene sets of differentially expressed genes were analysed in GSEA (Geneset Enrichment analysis) which also shows the related pathways. The significantly up-and downregulated genes (False Discovery Rate (FDR) < 0.05) and related pathways between CTL and AF/S chips and between CTL chip and COLIV insert are provided in Supplementary Tables 1, 2 and 3.

### cDNA and qPCR

For cDNA conversion, 500 ng of total RNA was reverse transcribed using oligo dT primers (Qiagen Benelux B.V., Venlo, The Netherlands) and M-MLV Polymerase (Thermo Fisher Scientific) at 42°C. All quantitative PCR (qPCR) reactions were performed in triplicate on a CFX-384 Real-Time PCR detection system (Bio-Rad Laboratories, Veenendaal, The Netherlands), using primers shown in Supplementary Table 6 and IQ SYBRGreen supermix (Bio-Rad). The relative standard curve method was used to calculate arbitrary gene expression using CFX-manager software (Bio-Rad). Two reference genes were included to calculate the normalized gene expression.

### CXCL8 ELISA

The protein encoded by CXCL8 (interleukin-8 or IL-8), was measured with use of the Recombinant Human IL-8/CXCL8 Protein ELISA kit of R&D (Abingdon, UK) according to manufacturer’s instructions and analyzed with use of 4-parameter logistic curve fitting in Prism 8 (version 8.1.1).

### Multiplex protein measurements

Cytokine and growth factor secretion was measured with the Bio-Plex Pro Human Cytokine Screening Panel (Bio-Rad #M500KCAF0Y), a 27-plex assay. Effluent culture medium was collected from the chips (collected in the Pod’s effluent reservoir) from cultures with 8 different donors. These samples were diluted 1:4 in assay buffer provided with the kit. Manufacturer’s instructions were followed to perform this assay, with minor adjustments to the protocol; one third of the proposed number of magnetic beads and detection antibodies was used. Plate readout was performed on a Luminex 100/200 System (Luminex, ‘s-Hertogenbosch, The Netherlands).

### IF staining and histology

At 14 day of ALI, cells in the Chip were fixed using 4% paraformaldehyde (PFA) solution. The PFA solution was pipetted into both channels and incubated for 20 min at RT, washed with PBS and stored in PBS at 4°C until staining. chips were subsequently removed from the carrier and cells were blocked and permeabilized using 0.5% (v/v) Triton-X 100 in PBS with 5% BSA for 60 min at RT. Chips were cut into 2 pieces using a razor blade and channels were filled with primary antibodies (Supplementary Table 7) diluted in the Triton/BSA buffer and incubated for 1h at RT. Samples were rinsed three times 5 min with PBS. Next, channels were filled with secondary antibodies diluted in Triton/BSA buffer which was left incubating for 1 h at RT, followed by a triple 5 min wash with PBS. 4′,6-Diamidino-2-phenylindole (DAPI, Sigma) was used to stain cell nuclei. Channels were next filled with prolong gold anti-fade (Thermo Scientific) and stored in the dark at 4 °C until imaging. Chips were imaged on Bellco coverslips, 25×75mm (Electron Microscopy Sciences, Hatfield, PA, SUA) with 0.13-0.17 mm thickness using a Leica DMi8 microscope equipped with an Andor Dragonfly 200 spinning disk confocal using a 10x objective (NA 0.30), 20x water objective (NA 0.50) or 40x water objective (NA 0.80).

### Cross-sectional imaging of thin chip slices

A new sectioning technique was developed to obtain cross-sectional slices of the chip for histology and immunofluorescence staining. Chips were fixed as described as above. The PDMS surrounding the chip channel was cut away as much as possible using a razor blade. Next, the chip channel was mounted on the specimen tube using glue (Pattex instant glue) of a semi-automated vibrating microtome (Compresstome^®^ VF-300-0Z, CMA, Kista, Sweden) and placed in the tube holder. Next, 150 µm thick slices were cut and collected in PBS-filled wells of a 12-well plate. Slices were stained in the well for classical H&E or immunofluorescence as described above. Slices were imaged on bellco coverslips as described above for the complete chip.

### IF Image processing and quantifications

#### Cell type quantification

Raw images were split into individual channels, median filtered, cropped and CLAHE-filtered^42^ to remove illumination artefacts, and auto-thresholded to retrieve the area fraction positive for each cell type marker. Semi-automated batch processing was conducted using ImageJ-Fiji^43^.

#### VANGL1 crescent quantification

Raw images were split into individual channels and cropped as needed to remove illumination artifacts. We used a Matlab script to de-noise the images with Gaussian, median, and Wiener filters and detected the VANGL1 crescents with Hessian based multiscale filtering (Supplementary Figure 6). The binarized crescents were skeletonized to yield lines of single pixel width and compute their total length. VANGL1 crescent density (D_C_) was defined as *D*_*C*_ *=* ∑ *L*_*i*_ */A*, where A is the area of the field of view in µm^2^, L_i_ is the length of VANGL1 crescent *i* in µm, and the sum ranges over all *i*=1…N crescents detected in the field of view. For each donor, final crescent density was reported as the average value from all fields of view.

### Measurements of CBF and MCC

#### Ciliary beat frequency measurement

CBF was measured using high-speed video microscopy followed by signal processing of the recordings as described previously in detail^10^. Briefly, to optimize optical imaging quality, inserts were washed for 10 minutes with PBS and the returned to ALI whereas Lung-Chips kept submerged in PBS or culture medium during imaging. Samples were imaged using a Leica DMi8 microscope equipped with phase-contrast objectives and a PCO Edge 4.2 high-speed camera with camera link interface that was operated from a PC desktop computer using the ImageJ-Fiji^43^ with Micro-Manager^44^. Per sample, 6 to 8 fields of view (166×166 µm, with ca. 3 pixel/µm spatial resolution) were recorded under Koehler illumination at a temporal resolution of 200 frames per second. Using a custom algorithm in Matlab (MathWorks), regions containing ciliary beat activity were detected automatically by computing the standard deviation of each pixel’s intensity over time, which reveals motion, and thresholding the resulting image using Otsu’s method to segment area with detected motion. Ciliary beat density was computed by dividing the number of pixels with detected motion by the total number of pixels in the image. In regions with ciliary beat, the dominant beat frequency was identified from the smoothed pixel intensity over time using Fast Fourier Transform (FFT). The ciliary beat frequencies and ciliary beat densities measured in all movies per sample were averaged for further statistical analysis.

#### Mucociliary clearance measurement

MCC was measured by adding fluorescent tracer particles to the top of samples, recording the movement of the tracers over time using video microscopy, and tracking the displacement of the particles in the resulting videos using specialized software to derive transport velocities. Briefly, the samples were washed and prepared as above before adding 30 µl of a 1:200 dilution of 1-µm sized fluorescent polystyrene microspheres (ThermoFisher Scientific) in PBS. Samples were imaged using a Leica DMi8 microscope equipped with a fluorescence light source and appropriate filters. Per sample, 3-6 fields of view (ca. 1200×1200 µm each) were recorded for 10 to 20 seconds at a temporal resolution of 30 frames per second. Using a batch-processing script in ImageJ-Fiji with the Trackmate plugin^45^, we tracked the bead displacement per time step to derive their trajectories and flow speed. To identify and filter out potential background flow, we used a supervised machine learning approach in Matlab to classify trajectories as either background or true signal. Briefly, we imaged bead movements in cell-free chips and inserts as well as in strong MCC flow. To generate the ground truth data, we labeled ca. 40,000 trajectories correctly as either true signal (MCC) or background flow. To characterize the trajectories, we measured the following metrics: mean speed, median speed, standard deviation of speed, path length, Euclidian distance covered, the ratio of Euclidian distance covered to path length (directness), mean flow angle, number of nodes, and maximal acceleration. Using the Matlab classification learner app, we trained a decision tree model with 5-fold cross-validation to identify true and background trajectories. We found that Euclidian distance, median speed, and directness below a threshold level were sufficient indicators to classify trajectories as background flow with an accuracy of more than 95%. This simple classification strategy was then used to automatically filter out the background flow in all data sets (Supplementary Figure 9). Matlab was used to further analyze the flow as described below.

#### MCC polar order parameter

The main direction of each bead trajectory was determined by the measuring the angle θ_*i*_ formed between the channel axis and the line between start point and end point of each bead trajectory (Figure 2d), where *i* ranges from 1 to the number of trajectories per sample. For each sample, an average unit vector 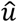 was determined from all measured angles: 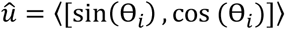, where the outer brackets indicate averaging. The PO was computed by dotting the average unit vector with the unit vector 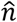 pointing towards the channel inlet: 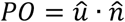. The PO hence ranges from -1 to 1, with -1 and 1 indicating perfect alignment of the mean flow vector towards the channel outlet or inlet, respectively. Intermediate values indicate partial alignment towards one the channel ends, i.e., negative values towards outlet and positive values towards inlet. A PO near 0 indicates isotropic flow or alignment of the mean flow vector towards the walls. Trajectory data from all movies per sample were pooled for the analysis. Therefore, alternating, perpendicular, and random flow without any bias would all be expected to result in an average PO of zero.

#### MCC coverage and speed

To measure MCC coverage and area-averaged speed, the trajectories were converted into a Eulerian velocity vector field *U(x, y) = u(x, y) î + v(x, y) î* by averaging the flow velocity component in x direction (*u*) and in y direction (*ν*) measured at each coordinate point over time (since multiple beads may pass over the same location) (Supplementary Fig. 11a left and center). The Matlab “alphaShape” function was used to generate coherent areas from these point measurements and estimate MCC coverage *C*_*MCC*_, i.e., the fraction of surface area with measurable MCC: 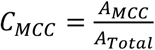 where *A*_*MCC*_ is the area with measurable MCC and *A*_*Total*_ is the total tissue surface area visible in the field of view (Supplementary Fig. 11a right). Area-averaged speed 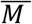 was computed by weighting the average flow speed in active regions by their relative surface fraction: 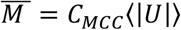, where | *U* | is the scalar field of flow magnitudes and the brackets indicate averaging. Flow data from all movies per sample were pooled for statistical analysis.

### Statistical methods

Statistical analysis was conducted using GraphPad Prism version 8 and 9 (GraphPad Software Inc). Differences were considered significant at p< 0.05. Statistical significance, statistical methods, and the number of donors and samples used are detailed in the figures and figure legends. Where number of donors is reported, this always refers to the number of different donors.

