## Supplemental Material for "Breathing on Chip: Dynamic flow and stretch tune cellular composition and accelerate mucociliary maturation of airway epithelium *in vitro*"

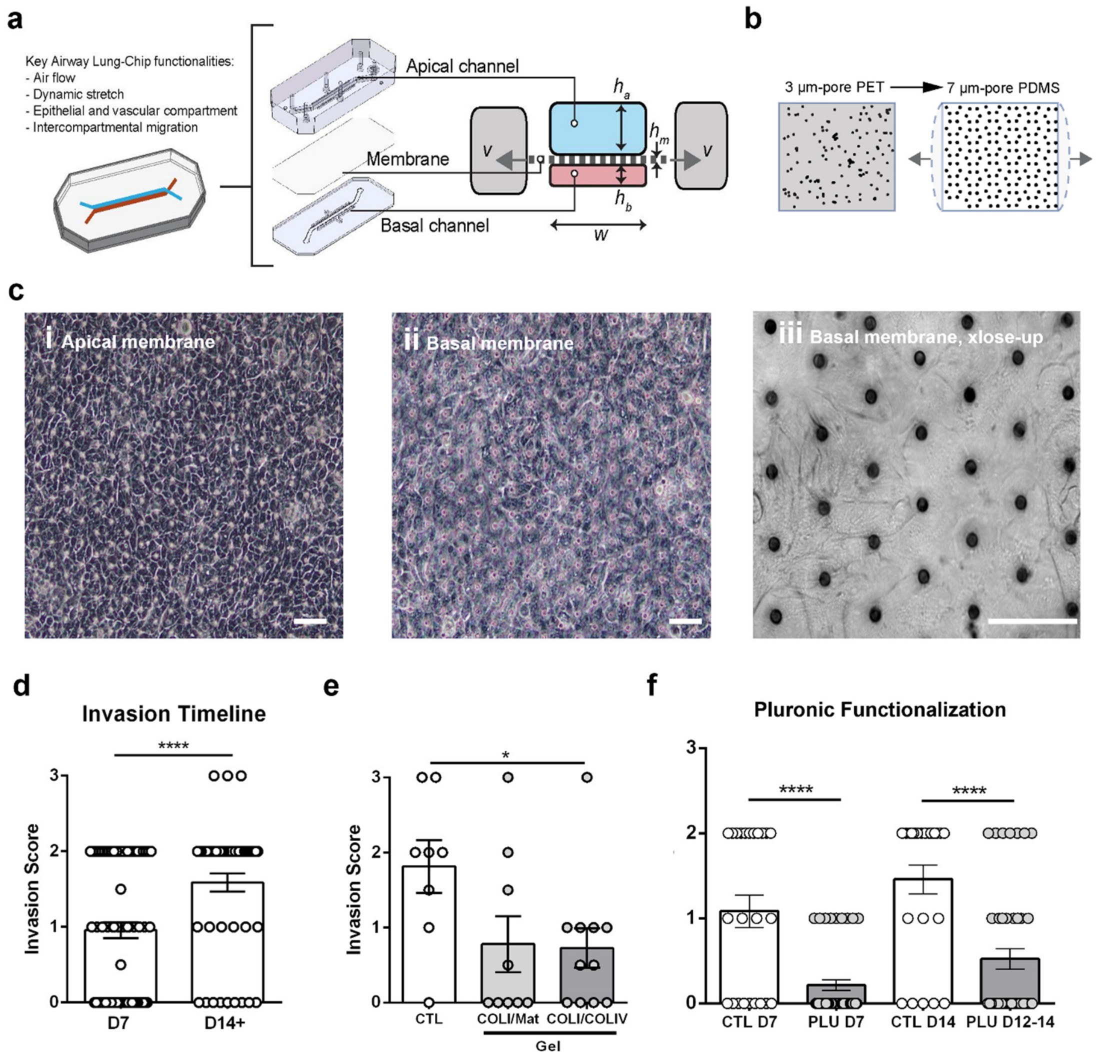

**Supplementary Figure 1: Benefits and challenges of the Chip S1 design for pHBEC culture.**

**a**, The Chip-S1 supports key functionalities for in vitro airway studies through the perfusable dual-compartment design and porous vacuum(v)-actuatable membrane. w, channel width = 1000 µm; ha, top channel height = 1000 µm; hb, bottom channel height = 200 µm; hm, membrane thickness = 25 µm. **b**, Previous airway chip studies commonly used a rigid and translucent PET membrane. To enable the current study, we switched to the Chip-S1's flexible PDMS membrane with optical transparency. **c**, Phase contrast microscopy of top of membrane showing pHBEC at ALI (left) and bottom of membrane showing a confluent layer of pHBEC that have migrated through the 7-µm pores to the bottom channel (center and right). Scalebars: 50 µm. **d**, Quantification of bottom channel invasion by hPBEC on day 7 and after day 14 of ALI using a score of 0 (no invasion) to 3 (basal channel fully lined). Data pooled from multiple media and membrane coating conditions. N=4 donors with 2-6 chips per donor. Each data point is from 1 chip. Columns and error bars represent mean +/- SEM, ****p<0.0001, Mann-Whitney test. **e**, Reduced invasion at day 14 ALI in chips with collagen I/Matrigel (COLI/Mat) or collagen I/collagen IV (COLI/COLIV) hydrogels. N=1 donor; each data point is from 1 chip. Data are depicted as mean +/- SEM. *p<0.05, ANOVA, followed by Dunnett's multiple comparison test. **f**,

Strongly reduced invasion in chips treated with Pluronic. N=3 donors, 12 chips per donor, each data point is from 1 chip. CTL data are the same as in Figure 1d. Columns and error bars represent mean +/- SEM. ****, p<0.001, Mann-Whitney test.

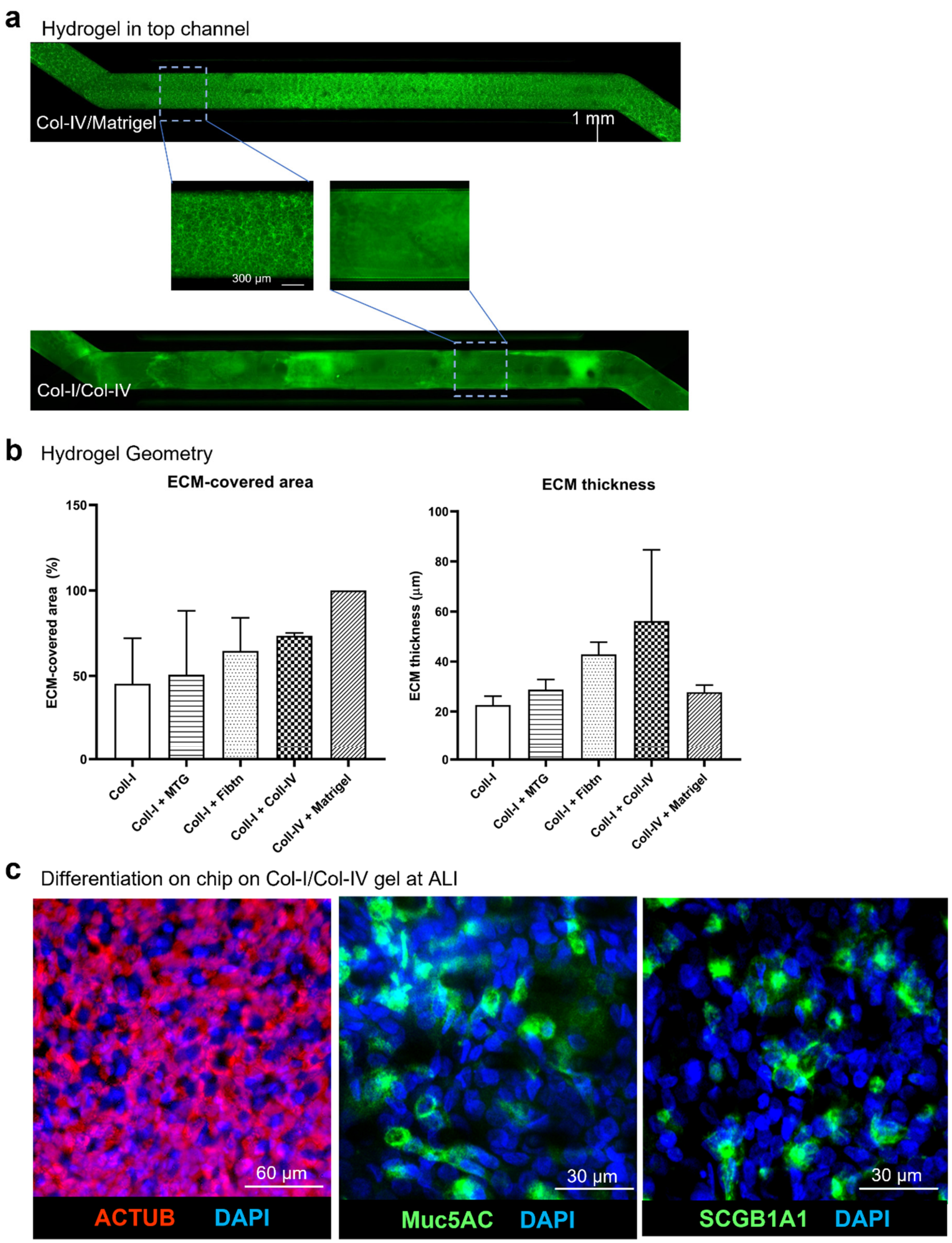

**Supplementary Figure 2: Hydrogel barrier supports pHBEC differentiation at ALI.**

**a**, Representative images of hydrogels (green) consisting of collagen I/Matrigel or collagen I/collagen IV lining the entire membrane in the top channel of the chip. **b**, Compared to other gel formulations, the gel compositions in (A) achieved superior percentage of membrane coverage (left) combined with robust gel thicknesses above 20 µm (right). Columns and error bars represent mean +/- SEM. **c**, Immunofluorescent (IF) staining of hydrogel chips at day 14 ALI shows presence of 3 major differentiated airway epithelial cell types: acetylated α-tubulin (ciliated cells), Muc5AC (goblet cells), SCGB1A1 (club cells). Nuclei are stained with DAPI.

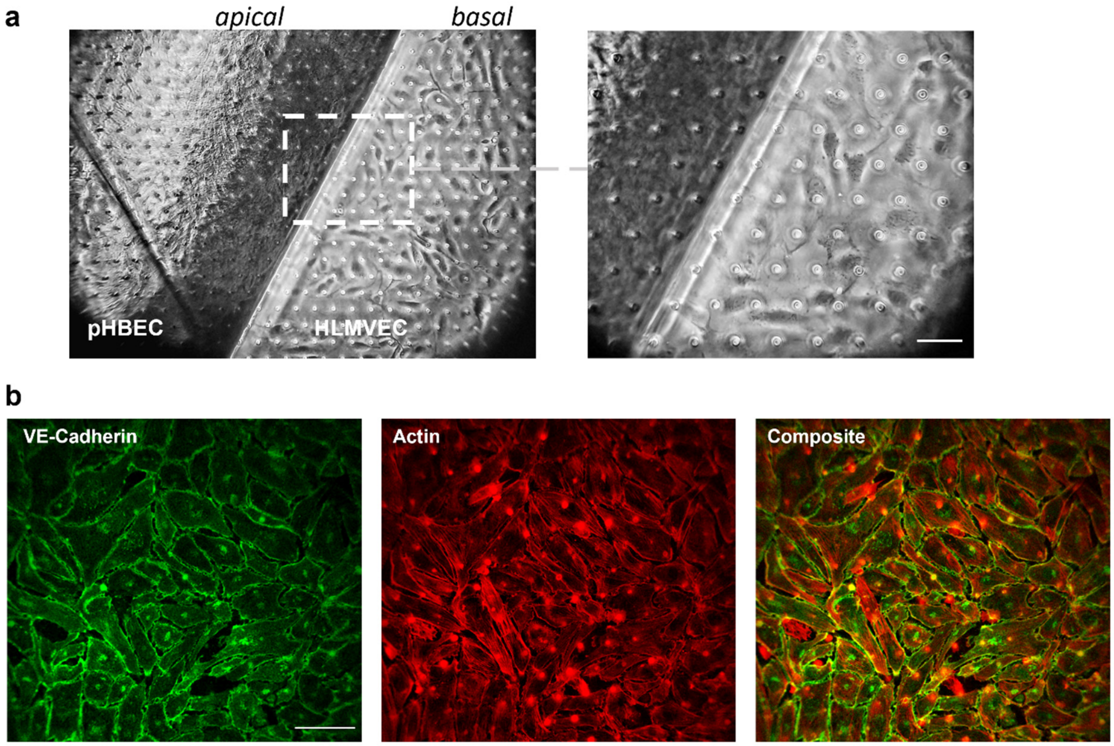

**Supplementary Figure 3: Endothelial co-culture on invasion-free airway epithelial chips.**

**a,** Left: Phase contrast image of co-culture of HPBEC in apical (top) channel and HLMVEC in basal (bottom) channel, on opposing sides of the PDMS membrane (shown here is junction where the channels diverge), 3h after seeding of the endothelial cells at day 12 ALI. Right: inset showing typical endothelial morphology and successful adhesion to the fibronectin-coated PDMS membrane. Scale bar: 50 µm. **b,** IF stain showing that HLMVEC lining the basal channel membrane express the endothelial-specific cell junction marker vascular endothelial (VE)-cadherin at day 14 ALI, i.e., 48h post-seeding of the endothelial cells. Note that the regularly spaced, circular structures are the auto-fluorescent membrane pores. Scale bar: 100 µm

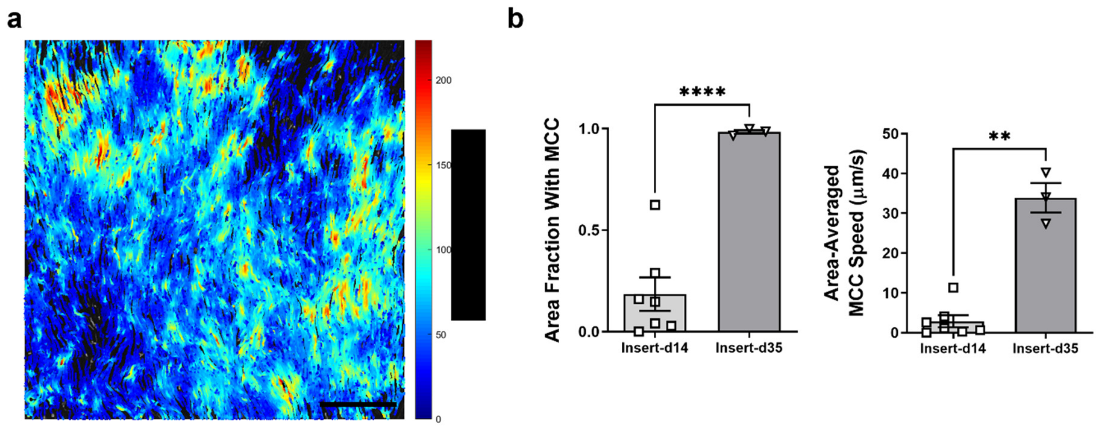

**Supplementary Figure 4: Mucociliary differentiation in inserts is improved by day 35 ALI.**
**a,** Example of fluorescent bead flow speeds in insert at day 35 ALI. Scale bar: 200 μm. **b**, Quantification of area fraction covered by MCC (left) and area-averaged MCC speed (right). Depicted are mean +/- SEM of 7 inserts at day 14 ALI from N=7 donors (one insert per donor) and 3 inserts from N=3 donors (one insert per donor) at day 35 ALI. **p<0.01, ****p<0.0001; two-tailed Welch's t-test.

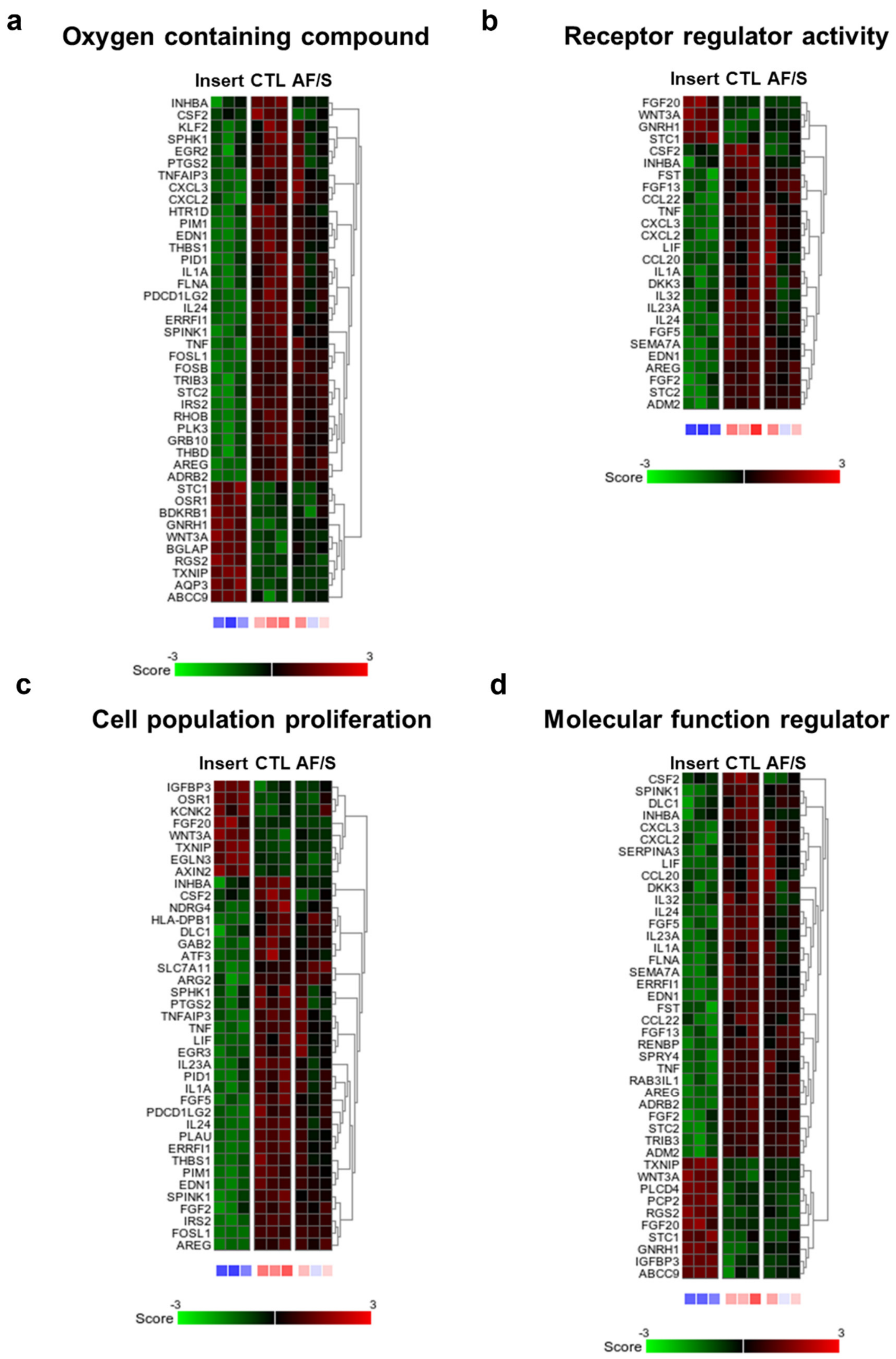

**Supplementary Figure 5: Changes in gene expression related to top pathways identified by comparison of insert with chip cultures.**

**a,** Heat maps displaying the Z score of DEGs related to the oxygen-containing compound pathway identified in pathway analysis of the DEGs identified between insert and chip cultures, which are depicted in Table 2. **b,** Heat maps displaying the Z score of DEGs related to the receptor regulator pathway. **c,** Heat maps displaying the Z score of DEGs related to the cell population proliferation pathway. **d,** Heat maps displaying the Z score of DEGs related to the molecular function regulator pathway. Expression was compared between insert, CTL chip and AF/S chip cultures with N=3 donors that are paired (one donor per column).

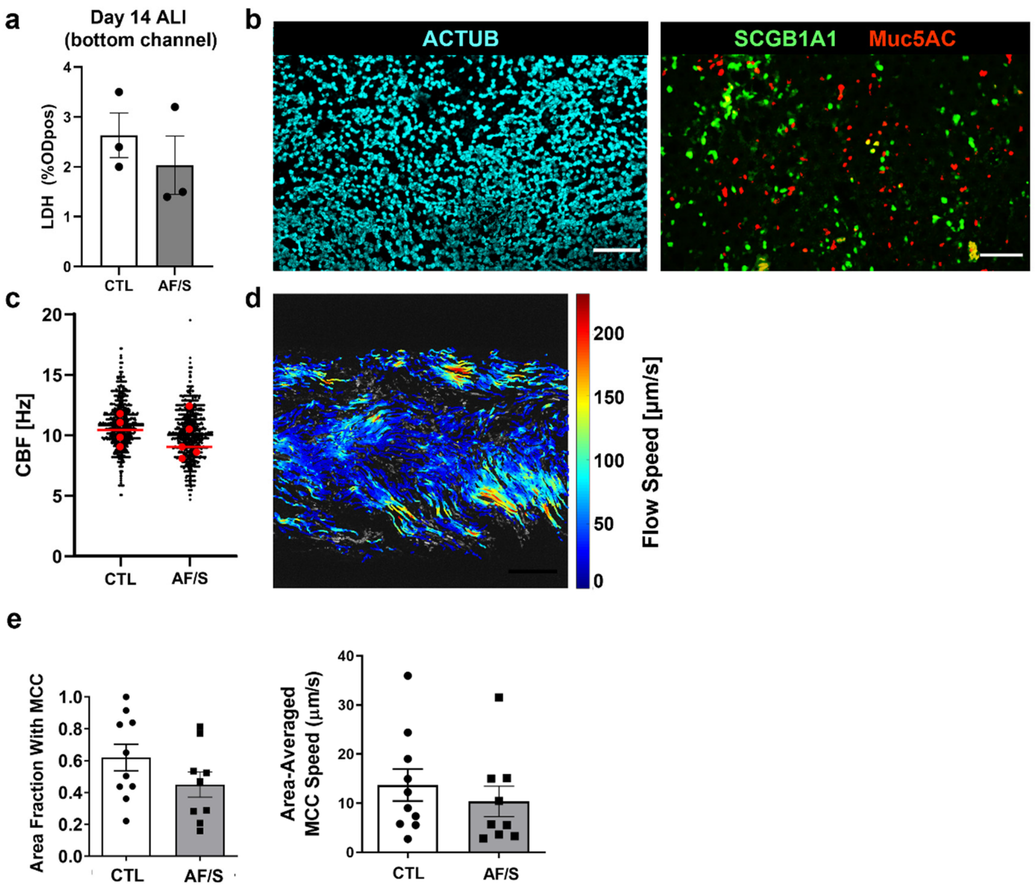

**Supplementary Figure 6: Mucociliary differentiation is comparable between CTL and AF/S chips.**
**a,** LDH analysis on medium collected from the bottom channel reservoir after 24 h of flowing between day 13 and 14 of ALI. Data are from N=3 different donors, 1 chip each. **b,** Representative IF staining of club (SCGB1A1), goblet (muc5ac), and ciliated cells (acetylated alpha-tubulin, ACTUB) in AF/S chip Scalebar: 100 µm. **c,** CBF analysis. Data from 4 biological replicates per condition from N=2 donors each. Each black dot represents approximately 1 ciliated cell (CTL Chip: 2481; AF/S Chip: 2046). Red dots indicate individual means of biological replicates and line indicates their median. **d,** Example of fluorescent bead flow speeds in AF/S chip. Scale bar: 200 µm. **e**, Quantification of area fraction covered by MCC (left) and area-averaged MCC speed (right). Depicted are mean +/- SEM of 8 AF/S chips from N=8 donors (one chip per donor) and 9 CTL chips from N=9 donors (one chip per donor). *, $p<0.05$; **, $p<0.01$; two-tailed Welch's t-test.

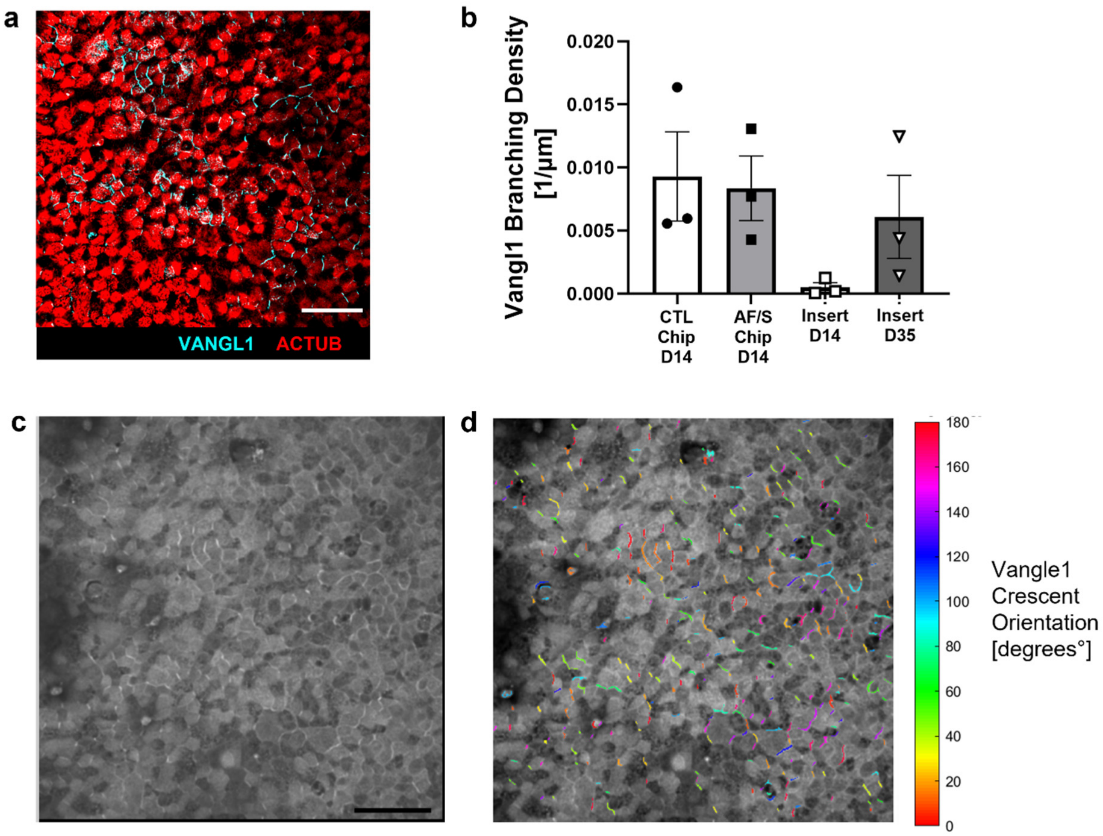

**Supplementary Figure 7: VANGL1 crescent density is comparable between chips at day 14 of ALI and inserts at day 35 of ALI.**

**a,** Representative staining of VANGL1 at day 35 ALI in same donor as in Fig.3 at day 14 ALI. Scale bar: 50 µm. **b,** Quantification of VANGL1 crescent density in chips at day 14 ALI compared to inserts at day 14 ALI and day 35 ALI. N=3 donors, one chip or insert each; each point is mean value from 1 insert or chip; columns and error bars represent mean +/- SEM. .**c,** Raw intensity image of VANGL1 IF stain. Scale bar: 50 µm. **d,** Detected VANGL1 crescents using Hessian matrix based multiscale analysis of curvature in Matlab.

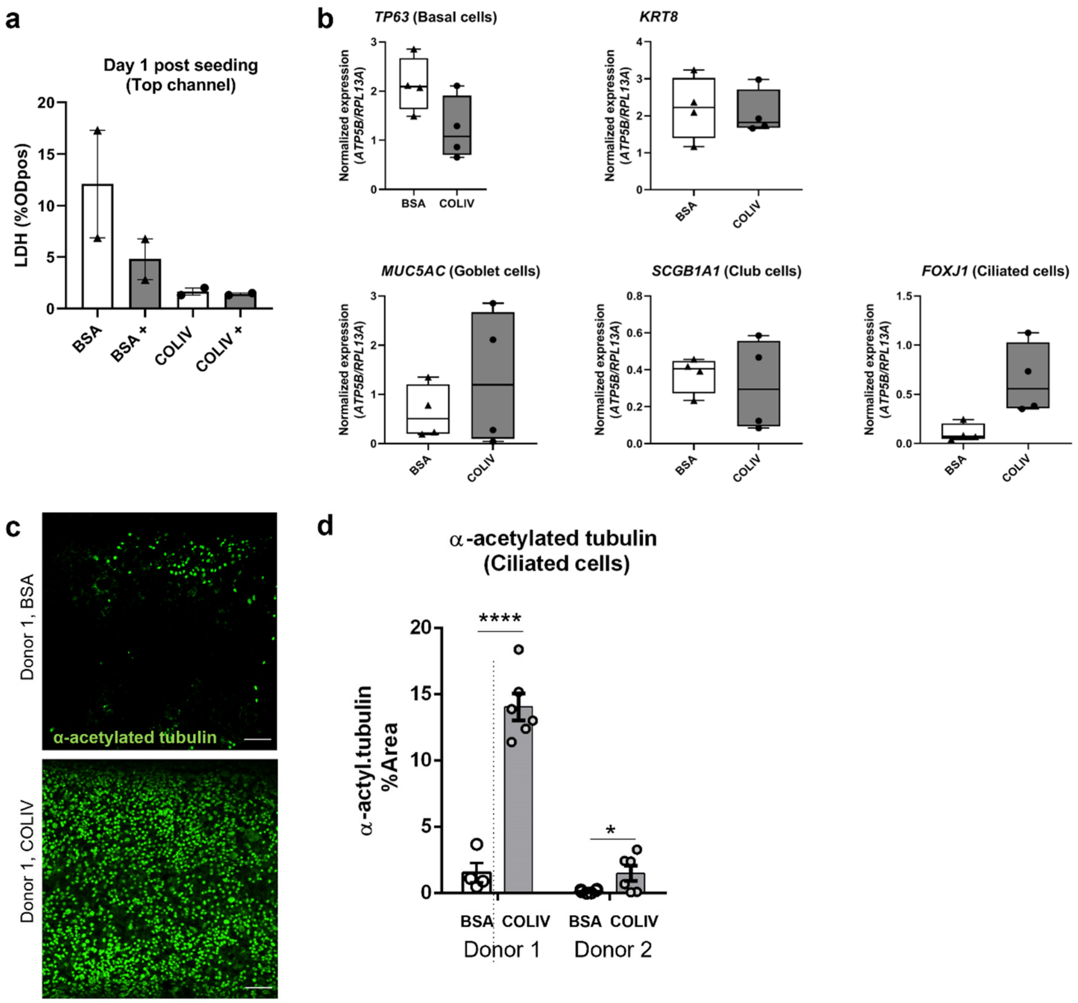

**Supplementary Figure 8: Beneficial effect of Collagen IV coating on pHBEC viability and differentiation on Chip.**

**a,** LDH analysis on medium collected from the top channel reservoir after 24 h of flowing, 24 h after seeding from chips coated with BSA, BSA and pluronic treatment (+) or collagen 4 (COLIV) with and without pluronic treatment. Data are from N=2 donors, 1 chip per donor. Data are depicted as mean +/- SEM. **b,** At 14 days post air-liquid interface (ALI) cells were lysed and RNA was isolated followed by cDNA synthesis to assess gene expression of *TP63* (basal cells), *KRT8* (intermediate cells), *MUC5AC* (goblet cells), *SCGB1A1* (club cells) and *FOXJ1* (ciliated cells). Triangles: BSA-coated chips, black circles: COLIV-coated chips, data are shown as target gene expression normalized for the geometric mean expression of the reference genes ATP synthase, H+ transporting, mitochondrial F1 complex, beta polypeptide (*ATP5B*), β2-microglobulin (B2M) and Ribosomal Protein L13a (*RPL13A*); N=4 different donors, one chip/donor. **c,** Representative IF stains of cilia in BSA- and COLIV- coated chips. **d,** Quantification of surface area fraction positive for cilia staining for N= 2 donors, 1 chip per donor. Each data point represents one field of view. Scalebar: 100 µm. Column and error bars represent mean +/- SEM. *p<0.05, ****p<0.0001, t-test with Holm-Sidak multiple comparison correction.

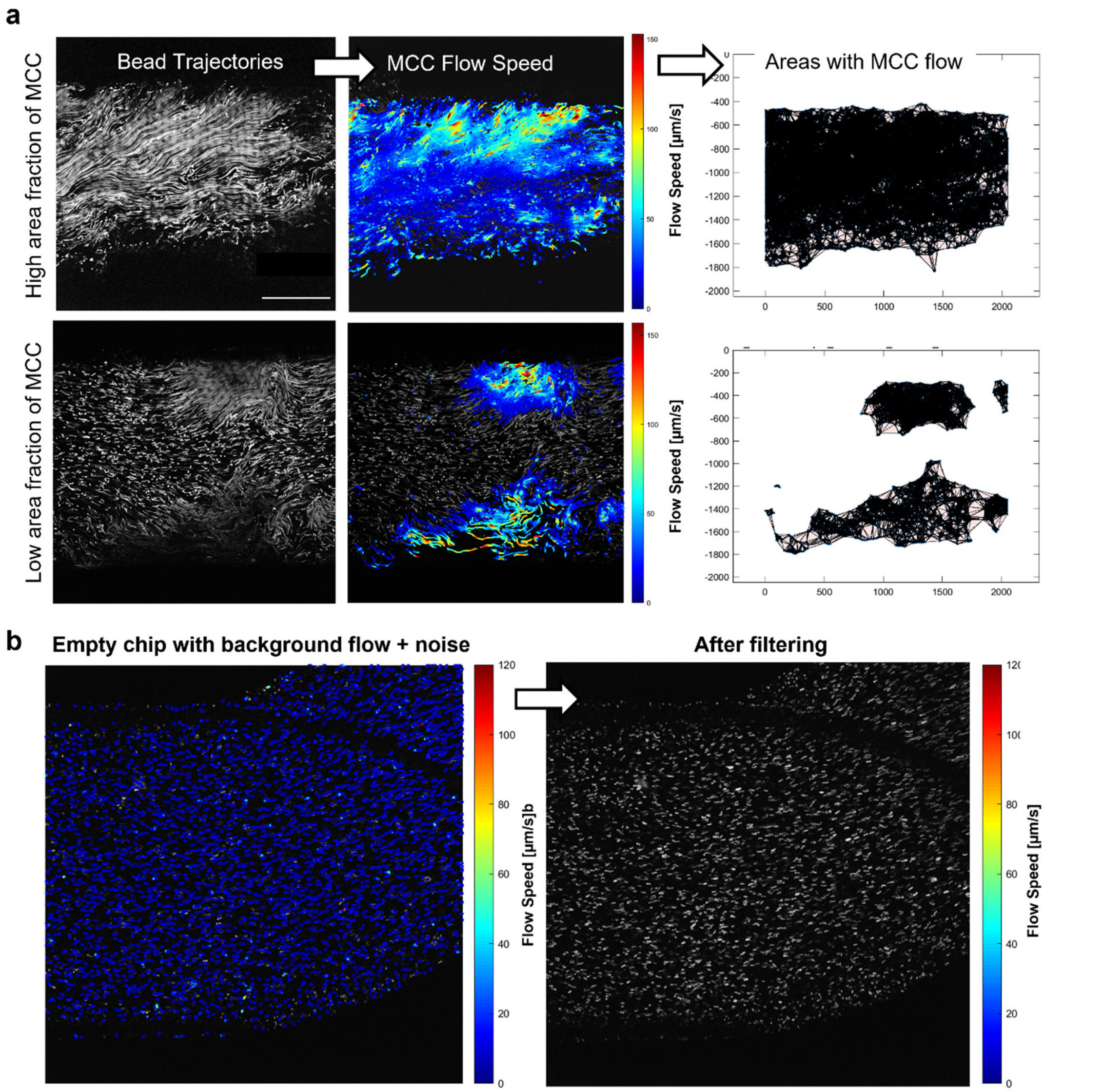

**Supplementary Figure 9: MCC quantification and noise removal.**

**a**, MCC flow speed along each bead trajectory is determined from bead displacement (left and center column), then averaged for each coordinate in order to get the Eulerian flow field that allows estimating the continuous areas with MCC flow (right column). Trajectories due to background flow or noise are filtered out based on trajectory speed, length, and directness, as seen in comparison of movies with high versus low fraction of MCC (top and bottom row). Scalebar 500 µm. **b,** Detected flow speeds in empty chip (left, colored lines overlaid on bead trajectories) are ignored (right) after filtering with parameters determined by machine learning approach.

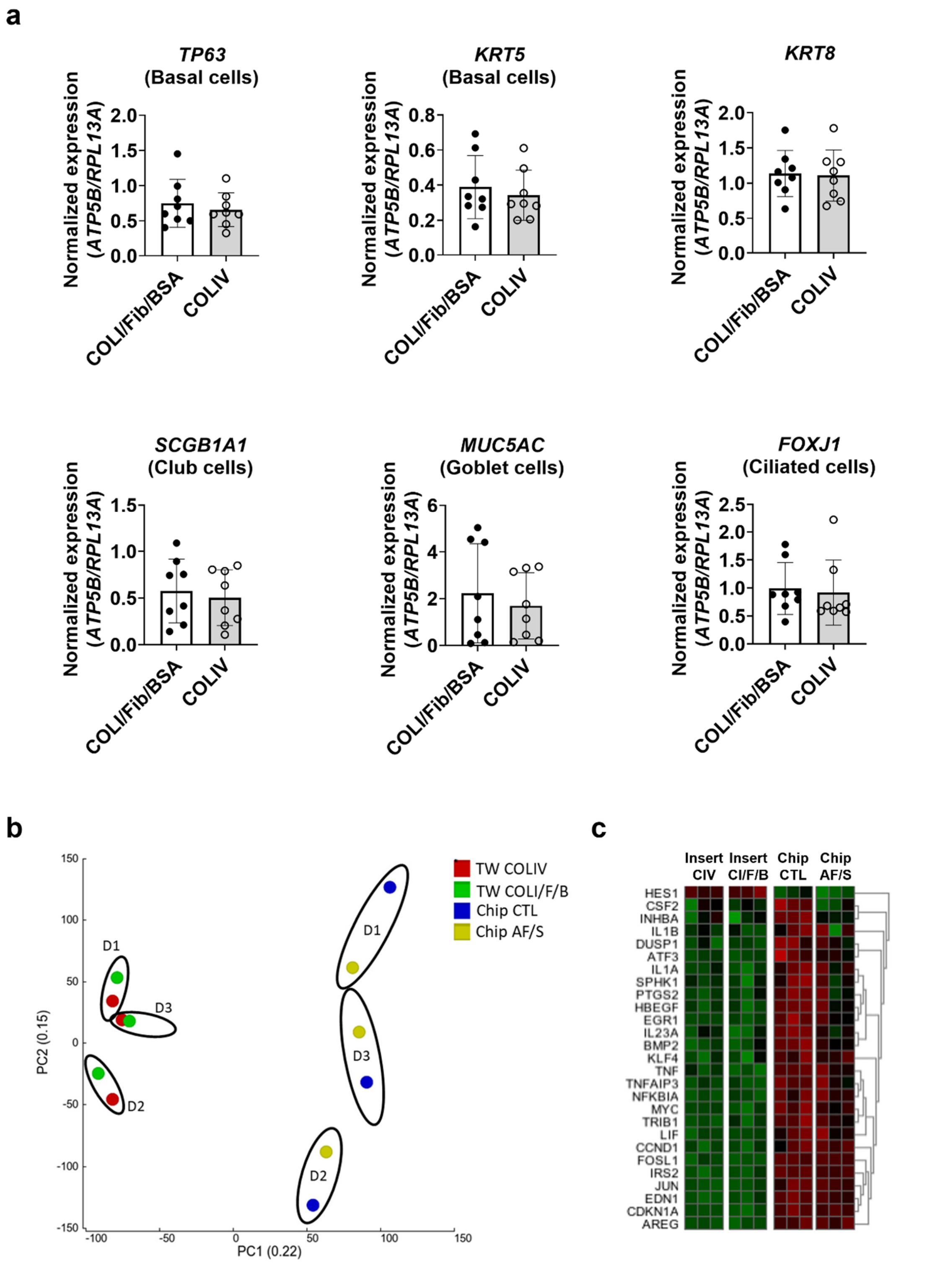

**Supplementary Figure 10: comparison of insert cultures with different coatings.**

**a,** Gene expression of *TP63* (basal cells), *KRT5* (basal cells), *KRT8* (differentiated non-basal cells), *SCGB1A1* (club cells), *MUC5AC* (goblet cells) and *FOXJ1* (ciliated cells) in collagen IV (COLIV)-coated or Collagen 1 (COLI), fibronectin and BSA-coated inserts at day 14 ALI. Data are shown as target gene expression normalized for the geometric mean expression of the reference genes *ATP5B* and *RPL13A*; N=8 donors, one chip or insert per donor. Data are depicted as mean +/- SEM. **b**, PCA on all transcriptomes showing distinct clusters of airway epithelial cells from N=3 donors in CTL chips, AF/S chips, or cell culture inserts coated with collagen IV or collagen I, fibronectin and BSA. **c,** Heat maps displaying the Z score of selected DEGs between insert, in CTL chips, AF/S chips, or cell culture inserts coated with collagen IV or collagen I, fibronectin and BSA, that were most frequently expressed in the top related pathways when comparing COLIV insert and CTL Chip with N=3 donors that are paired between insert, CTL chip and AF/S chip cultures (one donor per column).

**Supplementary Table 1.** Differentially expressed genes (*n*=182) between airway epithelial cells cultured on chip compared to cultures on inserts.

| S.No | ID | Base Mean | log2 Fold Change | q value | Gene Name |
|---|---|---|---|---|---|
| 1 | ENSG00000138166 | 2554.40 | 2.71 | 2.21E-56 | DUSP5 |
| 2 | ENSG00000175592 | 1178.69 | 2.54 | 9.51E-56 | FOSL1 |
| 3 | ENSG00000142871 | 5839.09 | 2.92 | 2.65E-42 | CYR61 |
| 4 | ENSG00000067082 | 6509.70 | 1.94 | 6.32E-39 | KLF6 |
| 5 | ENSG00000005108 | 316.95 | 3.50 | 8.53E-33 | THSD7A |
| 6 | ENSG00000173706 | 2357.66 | 1.86 | 1.21E-31 | HEG1 |
| 7 | ENSG00000116285 | 6509.04 | 1.74 | 6.55E-31 | ERRFI1 |
| 8 | ENSG00000112182 | 197.20 | 2.76 | 2.03E-26 | BACH2 |
| 9 | ENSG00000125740 | 214.15 | 2.80 | 6.3E-26 | FOSB |
| 10 | ENSG00000117289 | 26962.15 | -1.60 | 1.27E-25 | TXNIP |
| 11 | ENSG00000185950 | 2895.95 | 1.88 | 1.75E-25 | IRS2 |
| 12 | ENSG00000118503 | 5539.40 | 2.32 | 2.42E-24 | TNFAIP3 |
| 13 | ENSG00000122861 | 10509.12 | 1.56 | 2.89E-24 | PLAU |
| 14 | ENSG00000232810 | 72.60 | 3.45 | 5.27E-24 | TNF |
| 15 | ENSG00000078401 | 1044.29 | 3.02 | 6.7E-24 | EDN1 |
| 16 | ENSG00000135069 | 3214.49 | 2.41 | 8.52E-24 | PSAT1 |
| 17 | ENSG00000128965 | 829.91 | 1.87 | 5.34E-22 | CHAC1 |
| 18 | ENSG00000129521 | 1306.88 | -1.65 | 3.64E-19 | EGLN3 |
| 19 | ENSG00000137193 | 2099.09 | 1.82 | 3.67E-19 | PIM1 |
| 20 | ENSG00000169252 | 1196.70 | 1.53 | 8.47E-19 | ADRB2 |
| 21 | ENSG00000168003 | 10338.71 | 1.60 | 1.57E-18 | SLC3A2 |
| 22 | ENSG00000078081 | 885.06 | 2.10 | 1.52E-17 | LAMP3 |
| 23 | ENSG00000179148 | 78.37 | 3.03 | 4.5E-17 | ALOXE3 |
| 24 | ENSG00000137801 | 13490.22 | 2.15 | 1.03E-16 | THBS1 |
| 25 | ENSG00000151012 | 2458.37 | 1.85 | 1.91E-16 | SLC7A11 |
| 26 | ENSG00000143878 | 3339.03 | 1.71 | 2.19E-16 | RHOB |

| | | | | | |
|---|---|---|---|---|---|
| 27 | ENSG00000128591 | 111.52 | 2.83 | 2.2E-16 | FLNC |
| 28 | ENSG00000122254 | 314.28 | 3.08 | 4.69E-16 | HS3ST2 |
| 29 | ENSG00000187840 | 1340.47 | 1.53 | 4.98E-16 | EIF4EBP1 |
| 30 | ENSG00000101255 | 3540.07 | 2.90 | 3.88E-15 | TRIB3 |
| 31 | ENSG00000165272 | 27793.94 | -1.52 | 9.26E-15 | AQP3 |
| 32 | ENSG00000116741 | 526.16 | -1.74 | 1.02E-14 | RGS2 |
| 33 | ENSG00000139289 | 3564.85 | 1.57 | 1.22E-14 | PHLDA1 |
| 34 | ENSG00000168646 | 840.17 | -1.84 | 5.62E-14 | AXIN2 |
| 35 | ENSG00000221968 | 873.33 | 2.06 | 1.08E-13 | FADS3 |
| 36 | ENSG00000179388 | 298.44 | 2.27 | 1.79E-13 | EGR3 |
| 37 | ENSG00000212724 | 38.40 | 2.93 | 2.96E-13 | KRTAP2-3 |
| 38 | ENSG00000163734 | 705.93 | 2.24 | 3.99E-13 | CXCL3 |
| 39 | ENSG00000078018 | 1155.01 | 2.23 | 8.65E-13 | MAP2 |
| 40 | ENSG00000130598 | 266.37 | -1.83 | 1.07E-12 | TNNI2 |
| 41 | ENSG00000138675 | 106.24 | 2.27 | 1.78E-12 | FGF5 |
| 42 | ENSG00000102032 | 92.33 | 2.64 | 2.22E-12 | RENBP |
| 43 | ENSG00000103257 | 13838.64 | 1.87 | 2.79E-12 | SLC7A5 |
| 44 | ENSG00000109321 | 198.64 | 1.86 | 4.12E-12 | AREG |
| 45 | ENSG00000033327 | 513.55 | 1.64 | 1.03E-11 | GAB2 |
| 46 | ENSG00000173846 | 973.85 | 1.68 | 1.2E-11 | PLK3 |
| 47 | ENSG00000074416 | 1878.28 | 1.55 | 1.24E-11 | MGLL |
| 48 | ENSG00000163638 | 877.13 | 2.64 | 1.92E-11 | ADAMTS9 |
| 49 | ENSG00000153132 | 855.93 | 1.84 | 2.09E-11 | CLGN |
| 50 | ENSG00000196167 | 5387.56 | -1.51 | 4.03E-11 | COLCA1 |
| 51 | ENSG00000115556 | 112.75 | -1.87 | 6.15E-11 | PLCD4 |
| 52 | ENSG00000106070 | 1612.77 | 1.59 | 7.16E-11 | GRB10 |
| 53 | ENSG00000070669 | 3431.99 | 1.52 | 7.92E-11 | ASNS |
| 54 | ENSG00000128165 | 650.51 | 2.10 | 8.87E-11 | ADM2 |
| 55 | ENSG00000138623 | 1978.41 | 2.46 | 9.91E-11 | SEMA7A |
| 56 | ENSG00000079308 | 5281.32 | -1.60 | 1.81E-10 | TNS1 |
| 57 | ENSG00000178726 | 1449.10 | 1.75 | 2.06E-10 | THBD |
| 58 | ENSG00000130294 | 34.29 | 2.61 | 2.09E-10 | KIF1A |
| 59 | ENSG00000113739 | 2337.69 | 1.96 | 2.12E-10 | STC2 |
| 60 | ENSG00000233539 | 305.89 | -1.55 | 2.14E-10 | AC011294.3 |
| 61 | ENSG00000115008 | 807.50 | 2.22 | 2.51E-10 | IL1A |
| 62 | ENSG00000143867 | 402.76 | -1.63 | 6.89E-10 | OSR1 |
| 63 | ENSG00000073756 | 5561.02 | 2.07 | 1.1E-09 | PTGS2 |
| 64 | ENSG00000011422 | 1160.57 | 1.68 | 1.23E-09 | PLAUR |
| 65 | ENSG00000244242 | 2970.49 | -1.51 | 1.4E-09 | IFITM10 |
| 66 | ENSG00000167995 | 115.69 | 2.09 | 1.5E-09 | BEST1 |
| 67 | ENSG00000092621 | 3482.85 | 1.58 | 2.34E-09 | PHGDH |
| 68 | ENSG00000189410 | 241.52 | 2.07 | 2.43E-09 | SH2D5 |
| 69 | ENSG00000187678 | 136.97 | 1.70 | 2.57E-09 | SPRY4 |
| 70 | ENSG00000153446 | 366.64 | -1.62 | 2.82E-09 | C16orf89 |
| 71 | ENSG00000197446 | 5021.70 | -1.88 | 2.82E-09 | CYP2F1 |

| | | | | | |
|---|---|---|---|---|---|
| 72 | ENSG00000147437 | 77.08 | -1.98 | 2.99E-09 | GNRH1 |
| 73 | ENSG00000153823 | 168.52 | 2.25 | 3.52E-09 | PID1 |
| 74 | ENSG00000115112 | 22267.74 | -1.55 | 3.82E-09 | TFCP2L1 |
| 75 | ENSG00000178031 | 108.60 | -2.47 | 7.05E-09 | ADAMTSL1 |
| 76 | ENSG00000172548 | 154.30 | 2.31 | 9.9E-09 | NIPAL4 |
| 77 | ENSG00000110944 | 106.43 | 1.98 | 9.9E-09 | IL23A |
| 78 | ENSG00000111981 | 122.02 | 2.32 | 1.03E-08 | ULBP1 |
| 79 | ENSG00000136404 | 110.39 | 1.81 | 1.04E-08 | TM6SF1 |
| 80 | ENSG00000162892 | 18.15 | 2.45 | 1.05E-08 | IL24 |
| 81 | ENSG00000177614 | 215.91 | 1.88 | 1.75E-08 | PGBD5 |
| 82 | ENSG00000162772 | 2355.44 | 1.84 | 1.79E-08 | ATF3 |
| 83 | ENSG00000169429 | 11410.93 | 1.69 | 1.82E-08 | IL8 |
| 84 | ENSG00000008517 | 631.02 | 1.97 | 1.85E-08 | IL32 |
| 85 | ENSG00000156510 | 399.76 | 1.94 | 1.92E-08 | HKDC1 |
| 86 | ENSG00000196924 | 40824.80 | 1.53 | 2.1E-08 | FLNA |
| 87 | ENSG00000103034 | 410.59 | 2.02 | 2.38E-08 | NDRG4 |
| 88 | ENSG00000174788 | 31.96 | -2.18 | 2.67E-08 | PCP2 |
| 89 | ENSG00000198759 | 247.00 | -1.52 | 3.36E-08 | EGFL6 |
| 90 | ENSG00000128342 | 1554.74 | 1.61 | 3.44E-08 | LIF |
| 91 | ENSG00000076706 | 407.26 | 2.15 | 5.34E-08 | MCAM |
| 92 | ENSG00000105499 | 65.05 | 2.17 | 6.95E-08 | PLA2G4C |
| 93 | ENSG00000197408 | 343.49 | -1.86 | 7.39E-08 | CYP2B6 |
| 94 | ENSG00000146674 | 67497.38 | -1.51 | 8.89E-08 | IGFBP3 |
| 95 | ENSG00000060982 | 825.50 | 1.82 | 1.59E-07 | BCAT1 |
| 96 | ENSG00000154342 | 60.52 | -1.87 | 1.62E-07 | WNT3A |
| 97 | ENSG00000164266 | 41.20 | 2.26 | 1.66E-07 | SPINK1 |
| 98 | ENSG00000223865 | 402.46 | 1.57 | 1.66E-07 | HLA-DPB1 |
| 99 | ENSG00000233930 | 108.61 | 1.53 | 1.81E-07 | KRTAP5-AS1 |
| 100 | ENSG00000081041 | 1159.03 | 2.03 | 1.96E-07 | CXCL2 |
| 101 | ENSG00000026559 | 405.12 | 1.83 | 2.07E-07 | KCNG1 |
| 102 | ENSG00000153294 | 1127.03 | 1.58 | 2.86E-07 | GPR115 |
| 103 | ENSG00000210195 | 105.15 | 1.56 | 3.86E-07 | MT-TT |
| 104 | ENSG00000206538 | 697.11 | 1.90 | 4.57E-07 | VGLL3 |
| 105 | ENSG00000127561 | 68.27 | 1.58 | 4.78E-07 | SYNGR3 |
| 106 | ENSG00000183837 | 39.15 | -1.83 | 5.82E-07 | PNMA3 |
| 107 | ENSG00000172602 | 470.02 | 1.78 | 9.54E-07 | RND1 |
| 108 | ENSG00000136010 | 473.92 | 2.02 | 1.5E-06 | ALDH1L2 |
| 109 | ENSG00000165868 | 150.33 | 1.54 | 1.52E-06 | HSPA12A |
| 110 | ENSG00000072041 | 215.36 | 1.85 | 1.71E-06 | SLC6A15 |
| 111 | ENSG00000164400 | 58.04 | 2.03 | 1.89E-06 | CSF2 |
| 112 | ENSG00000069431 | 770.80 | -1.82 | 2.32E-06 | ABCC9 |
| 113 | ENSG00000128422 | 9823.81 | 1.83 | 2.63E-06 | KRT17 |
| 114 | ENSG00000167994 | 20.90 | 2.07 | 2.8E-06 | RAB3IL1 |
| 115 | ENSG00000100739 | 41.71 | -1.67 | 2.88E-06 | BDKRB1 |
| 116 | ENSG00000197646 | 29.46 | 2.00 | 3.07E-06 | PDCD1LG2 |

| | | | | | |
|---|---|---|---|---|---|
| 117 | ENSG00000081181 | 337.51 | 1.61 | 4.73E-06 | ARG2 |
| 118 | ENSG00000134363 | 545.84 | 1.52 | 4.84E-06 | FST |
| 119 | ENSG00000162576 | 40.16 | 2.02 | 5.39E-06 | MXRA8 |
| 120 | ENSG00000122786 | 3998.71 | 1.54 | 5.71E-06 | CALD1 |
| 121 | ENSG00000130775 | 159.39 | 1.73 | 8E-06 | THEMIS2 |
| 122 | ENSG00000049323 | 1263.59 | 1.54 | 8.16E-06 | LTBP1 |
| 123 | ENSG00000249601 | 50.20 | -1.97 | 8.43E-06 | CTB-27N1.1 |
| 124 | ENSG00000112149 | 47.89 | 1.85 | 9.63E-06 | CD83 |
| 125 | ENSG00000179546 | 29.07 | 1.95 | 1.32E-05 | HTR1D |
| 126 | ENSG00000122641 | 472.75 | 1.80 | 1.63E-05 | INHBA |
| 127 | ENSG00000101335 | 1440.40 | 1.79 | 1.67E-05 | MYL9 |
| 128 | ENSG00000101670 | 637.25 | 1.57 | 1.73E-05 | LIPG |
| 129 | ENSG00000162614 | 144.05 | 1.81 | 1.83E-05 | NEXN |
| 130 | ENSG00000119938 | 147.15 | -1.52 | 1.97E-05 | PPP1R3C |
| 131 | ENSG00000128655 | 755.38 | -1.55 | 1.98E-05 | PDE11A |
| 132 | ENSG00000269962 | 28.06 | 1.85 | 2.25E-05 | RP13-238F13.5 |
| 133 | ENSG00000108551 | 139.51 | 1.87 | 2.43E-05 | RASD1 |
| 134 | ENSG00000152463 | 115.11 | 1.83 | 2.56E-05 | OLAH |
| 135 | ENSG00000135960 | 73.81 | -1.64 | 2.92E-05 | EDAR |
| 136 | ENSG00000158014 | 77.78 | 1.81 | 3.19E-05 | SLC30A2 |
| 137 | ENSG00000159167 | 556.99 | -1.56 | 3.42E-05 | STC1 |
| 138 | ENSG00000167767 | 4907.05 | 1.70 | 3.85E-05 | KRT80 |
| 139 | ENSG00000139278 | 774.67 | 1.68 | 3.89E-05 | GLIPR1 |
| 140 | ENSG00000184908 | 44.61 | -1.85 | 4.14E-05 | CLCNKB |
| 141 | ENSG00000166073 | 249.18 | 1.64 | 5.05E-05 | GPR176 |
| 142 | ENSG00000138685 | 170.59 | 1.81 | 5.45E-05 | FGF2 |
| 143 | ENSG00000123405 | 41.29 | -1.67 | 5.47E-05 | NFE2 |
| 144 | ENSG00000176170 | 1093.97 | 1.50 | 5.86E-05 | SPHK1 |
| 145 | ENSG00000251323 | 27.71 | 1.61 | 5.87E-05 | RP11-452H21.4 |
| 146 | ENSG00000139211 | 1824.88 | 1.60 | 6.25E-05 | AMIGO2 |
| 147 | ENSG00000101210 | 115.19 | 1.70 | 6.32E-05 | EEF1A2 |
| 148 | ENSG00000129682 | 77.01 | 1.66 | 6.32E-05 | FGF13 |
| 149 | ENSG00000169035 | 747.43 | -1.55 | 6.75E-05 | KLK7 |
| 150 | ENSG00000115009 | 852.75 | 1.60 | 6.77E-05 | CCL20 |
| 151 | ENSG00000168269 | 49.99 | -1.78 | 7.92E-05 | FOXI1 |
| 152 | ENSG00000127528 | 143.27 | 1.69 | 8.24E-05 | KLF2 |
| 153 | ENSG00000050165 | 1027.22 | 1.58 | 8.25E-05 | DKK3 |
| 154 | ENSG00000203632 | 33.22 | -1.55 | 0.000101 | AC007690.1 |
| 155 | ENSG00000196136 | 1211.05 | 1.51 | 0.000111 | SERPINA3 |
| 156 | ENSG00000146592 | 162.97 | 1.67 | 0.000119 | CREB5 |
| 157 | ENSG00000102962 | 63.74 | 1.62 | 0.00013 | CCL22 |
| 158 | ENSG00000242252 | 20.07 | -1.67 | 0.000143 | BGLAP |
| 159 | ENSG00000272666 | 34.66 | 1.73 | 0.00015 | CTA-384D8.35 |
| 160 | ENSG00000134668 | 769.22 | 1.54 | 0.000155 | SPOCD1 |
| 161 | ENSG00000203727 | 42.47 | 1.64 | 0.000185 | SAMD5 |

| | | | | | |
|---|---|---|---|---|---|
| 162 | ENSG00000268852 | 12.70 | 1.71 | 0.000199 | AC132872.2 |
| 163 | ENSG00000255400 | 13.20 | 1.66 | 0.000215 | RP13-631K18.5 |
| 164 | ENSG00000188175 | 28.09 | -1.70 | 0.000221 | HEPACAM2 |
| 165 | ENSG00000260953 | 11.85 | 1.68 | 0.000236 | RP11-426C22.6 |
| 166 | ENSG00000182168 | 40.46 | -1.62 | 0.000239 | UNC5C |
| 167 | ENSG00000109758 | 23.27 | 1.68 | 0.000253 | HGFAC |
| 168 | ENSG00000087128 | 57.30 | -1.55 | 0.000319 | TMPRSS11E |
| 169 | ENSG00000269959 | 11.58 | -1.64 | 0.000391 | SPACA6P-AS |
| 170 | ENSG00000139209 | 28.34 | 1.64 | 0.0004 | SLC38A4 |
| 171 | ENSG00000164741 | 282.16 | 1.53 | 0.000471 | DLC1 |
| 172 | ENSG00000078579 | 12.20 | -1.60 | 0.000488 | FGF20 |
| 173 | ENSG00000184551 | 30.21 | 1.57 | 0.000512 | AC132872.1 |
| 174 | ENSG00000124191 | 97.10 | 1.59 | 0.000653 | TOX2 |
| 175 | ENSG00000178150 | 39.68 | 1.54 | 0.000696 | ZNF114 |
| 176 | ENSG00000260604 | 286.44 | 1.52 | 0.000723 | RP1-140K8.5 |
| 177 | ENSG00000082482 | 23.00 | -1.58 | 0.000743 | KCNK2 |
| 178 | ENSG00000163395 | 16.45 | 1.57 | 0.000783 | IGFN1 |
| 179 | ENSG00000122877 | 99.60 | 1.50 | 0.001139 | EGR2 |
| 180 | ENSG00000149591 | 4655.50 | 1.52 | 0.001196 | TAGLN |
| 181 | ENSG00000081138 | 10.55 | -1.52 | 0.001329 | CDH7 |
| 182 | ENSG00000067445 | 20.64 | -1.50 | 0.001395 | TRO |

**Supplementary Table 2**. Details of top 10 pathways associated with DEGs between cultures on inserts and control chips.

| S.No | Pathway description [# genes] | Pathway Description | #DEGs | ID | p value | q value |
|---|---|---|---|---|---|---|
| 1 | GOBP_RESPONSE_TO_OXYGEN_CONTAINING_COMPOUND [1653] | Any process that results in a change in state or activity of a cell or an organism (in terms of movement, secretion, enzyme production, gene expression, etc.) as a result of an oxygen-containing compound stimulus. [GOC:pr, GOC:TermGenie] | 43 | GO:1901700 | 2.76 $e^{-22}$ | 2.87 $e^{-18}$ |
| 2 | GOMF_SIGNALING_RECEPTOR_REGULATOR_ACTIVITY [543] | Binds to and modulates the activity of a receptor. [GOC:ceb] | 27 | GO:0030545 | 4.15 $e^{-21}$ | 2.16 $e^{-17}$ |
| 3 | GOBP_TISSUE_DEVELOPMENT [1916] | The process whose specific outcome is the progression of a tissue over time, from its formation to the mature structure. [ISBN:0471245208] | 44 | GO:0009888 | 1.03 $e^{-20}$ | 3.56 $e^{-17}$ |

| | | | | | | |
|---|---|---|---|---|---|---|
| 4 | GOBP_REGULATION_OF_CELL_POPULATION_PROLIFERATION [1745] | Any process that modulates the frequency, rate or extent of cell proliferation. [GOC:jl] | 41 | GO:0042127 | 1.23 $e^{-19}$ | 3.12 $e^{-16}$ |
| 5 | GOMF_MOLECULAR_FUNCTION_REGULATOR [1953] | A molecular function regulator regulates the activity of its target via non-covalent binding that does not result in covalent modification to the target. Examples of molecular function regulators include regulatory subunits of multimeric enzymes and channels. Mechanisms of regulation include allosteric changes in the target and competitive inhibition. [GOC:dos, GOC:pt] | 43 | GO:0098772 | 1.5 $e^{-19}$ | 3.12 $e^{-16}$ |
| 6 | GOBP_LOCOMOTION [1921] | Self-propelled movement of a cell or organism from one location to another. [GOC:dgh] | 42 | GO:0040011 | 5.59 $e^{-19}$ | 9.7 $e^{-16}$ |
| 7 | GOMF_SIGNALING_RECEPTOR_BINDING [1550] | Binding to one or more specific sites on a receptor molecule, a macromolecule that undergoes combination with a hormone, neurotransmitter, drug or intracellular messenger to initiate a change in cell function. [GOC:bf, GOC:ceb, ISBN:0198506732] | 38 | GO:0005102 | 8.29 $e^{-19}$ | 1.23 $e^{-15}$ |
| 8 | GOBP_REGULATION_OF_MULTICELLULAR_ORGANGANISMAL_DEVELOPMENT [1394] | Any process that modulates the frequency, rate or extent of multicellular organismal development. [GOC:obol] | 36 | GO:2000026 | 1.53 $e^{-18}$ | 1.99 $e^{-15}$ |
| 9 | GOBP_REGULATION_OF_CELL_DEATH [1643] | Any process that modulates the rate or frequency of cell death. Cell death is the specific activation or halting of processes within a cell so that its vital functions markedly cease, rather than simply deteriorating gradually over time, which culminates in cell death. [GOC:dph, GOC:tb] | 38 | GO:0010941 | 5.73 $e^{-18}$ | 6.62 $e^{-15}$ |
| 10 | GOBP_CELL_MIGRATION [1556] | The controlled self-propelled movement of a cell from one site to a destination guided by molecular cues. Cell migration is a central process in the development and maintenance of multicellular organisms. [GOC:cjm, GOC:dph, GOC:ems, | 37 | GO:0016477 | 6.92 $e^{-18}$ | 7.19 $e^{-15}$ |

| | | |
|---|---|---|
| | | GOC:pf, Wikipedia:Cell_migration] |

**Supplementary Table 3.** Differentially expressed genes (*n*=20) between airway epithelial cells cultured in airflow and stretch (AF/S)-exposed chips and control chip cultures.

| S.No | ID | Base Mean | log2 Fold Change | q value | Gene Name |
|---|---|---|---|---|---|
| 1 | ENSG00000122641 | 523.795 | -1.587 | 1.38E-19 | INHBA |
| 2 | ENSG00000177614 | 235.737 | -1.124 | 3.50E-06 | PGBD5 |
| 3 | ENSG00000100311 | 564.862 | -0.993 | 0.000189 | PDGFB |
| 4 | ENSG00000212724 | 46.661 | -0.891 | 0.001414 | KRTAP2-3 |
| 5 | ENSG00000179362 | 59.103 | -0.851 | 0.004539 | HMGN2P46 |
| 6 | ENSG00000084731 | 1352.044 | -0.737 | 0.018856 | KIF3C |
| 7 | ENSG00000163216 | 29.640 | -0.705 | 0.033372 | SPRR2D |
| 8 | ENSG00000139926 | 3125.493 | -0.684 | 0.005365 | FRMD6 |
| 9 | ENSG00000125845 | 1002.406 | -0.669 | 0.034607 | BMP2 |
| 10 | ENSG00000206538 | 959.505 | -0.666 | 0.026932 | VGLL3 |
| 11 | ENSG00000173281 | 1044.277 | -0.626 | 0.047517 | PPP1R3B |
| 12 | ENSG00000180573 | 2059.068 | -0.621 | 0.000833 | HIST1H2AC |
| 13 | ENSG00000167535 | 1438.138 | -0.568 | 0.017703 | CACNB3 |
| 14 | ENSG00000105855 | 4780.501 | -0.482 | 0.025469 | ITGB8 |
| 15 | ENSG00000185650 | 13894.804 | -0.466 | 0.000771 | ZFP36L1 |
| 16 | ENSG00000151726 | 9698.947 | 0.427 | 0.026932 | ACSL1 |
| 17 | ENSG00000144908 | 2473.898 | 0.529 | 0.026932 | ALDH1L1 |
| 18 | ENSG00000168497 | 1020.886 | 0.576 | 0.004539 | SDPR |
| 19 | ENSG00000060762 | 1069.072 | 0.588 | 6.24E-05 | MPC1 |
| 20 | ENSG00000119711 | 731.348 | 0.622 | 0.000724 | ALDH6A1 |

**Supplementary Table 4.** Donor characteristics.

| | |
|---|---|
| **Number of donors** | 13 |
| **Male/Female** | 6/6* |
| **Age (years) mean [SD]** | 61.7 [9.4]* |
| **BMI mean [SD]** | 26.0 [4.7]* |
| **Smoking Status (non-/ex-/smokers)** | 5/5/1** |

Abbreviations: SD (standard deviation); BMI (body mass index)

*Data available from 12 donors; ** Data available from 11 donors

**Supplementary Table 5.** Media/inhibitor strategies for reducing epithelial migration to bottom channel.

| | |
|---|---|
| **Media types** | |
| PneumaCult ALI | StemCell 05001 |
| B/D complete using supplements without hydrocortisone | ScienCell custom order |
| PneumaCult ALI w/o hydrocortisone | StemCell 05001 |
| Vertex ALI medium | Neuberger et al, 2011[1] |
| Airway organoid medium | Sachs et al, 2019[2] |
| **Cell culture compounds** | |
| Retinoic acid | 15; 150; 1500 ng/ml; Sigma R2625 |
| Dexamethasone | 0.1; 1; 10 µM; Sigma D4902 |
| **Inhibitors** | |
| *AG-1478*<br><br>Epidermal growth factor receptor (EGFR) tyrosine kinase inhibitor | 30; 150; 300 nM; Sigma T4182 |
| *PD098059*<br><br>Mitogen-Activated Protein Kinase inhibitor | 10 µM; Sigma P215 |
| *Y-27632*<br><br>Rho-associated, coiled-coil containing protein kinase (ROCK) inhibitor | 5 µM; Cayman Chemical 10005583 |
| *GM6001*<br><br>Matrix metalloprotease (MMP) inhibitor | 0.2; 0.5; 25 nM; Sigma CC1000 |
| *SB202190 + A83-01*<br><br>P38 MAPK inhibitor + TGFβ kinase/activin receptor-like kinase (ALK 5) inhibitor | 500 nM; 500 nM; Sigma S7067; Tocris 2939 |
| *SB202190 + A83-01 + noggin* | 500 nM; 500 nM; 100 ng/ml Sigma S7067; Tocris 2939; Peprotech 120-10c |

| |
|---|
| P38 MAPK inhibitor + TGFβ ALK inhibitor + Bone Morphogenetic Protein (BMP) Inhibitor |

**Supplementary Table 6.** Primer sequences.

| Gene | forward sequence (5' to 3') | reverse sequence (5' to 3') |
|---|---|---|
| *ATP5B* | TCACCCAGGCTGGTTCAGA | AGTGGCCAGGGTAGGCTGAT |
| *RPL13A* | AAGGTGGTGGTCGTACGCTGTG | CGGGAAGGGTTGGTGTTCATCC |
| *KRT5* | AGGAGTTGGACCAGTCAACAT | TGGAGTAGTAGCTTCCACTGC |
| *TP63* | CCACCTGGACGTATTCCACTG | TCGAATCAAATGACTAGGAGGGG |
| *KRT8* | TCCTCAGGCAGCTATATGAAGAG | GGTTGGCAATATCCTCGTACTGT |
| *FOXJ1* | GGAGGGGACGTAAATCCCTA | TTGGTCCCAGTAGTTCCAGC |
| *SCGB1A1* | ACATGAGGGAGGCAGGGGCTC | ACTCAAAGCATGGCAGCGGCA |
| *MUC5AC* | CCTTCGACGGACAGAGCTAC | TCTCGGTGACAACACGAAAG |
| *RSPH4A* | GAAGGGACGTGAGCTATAACAAC | GCAGGTAAGCCTTAGCATTCTGA |
| *DNAH11* | CAACAGCTTACCTTTCACCTGA | TTCTTCCCATAAAGTAGCTTGCC |
| *VANGL1* | CCGATCCTGTGGAGGGATGA | AAACACCCGTGGCATGTCA |
| *PTGS2* | TAAGTGCGATTGTACCCGGAC | TTTGTAGCCATAGTCAGCATTGT |
| *MMP9* | ACCTCGAACTTTGACAGCGAC | GAGGAATGATCTAAGCCCAGC |
| *FN1* | TGGAGGAAGCCGAGGTTT | CAGCGGTTTGCGATGGTA |

**Supplementary Table 7.** Antibodies used for confocal imaging.

| Antibody | Supplier | Catalog # | species | Antibody dilution |
|---|---|---|---|---|
| CK-8 | Millipore | MABT329 | rat | 1:100 |
| p63 | Abcam | ab124762 | rabbit | 1:100 |
| Mucin 5AC | Labvision Neomarkers | MS-145-P1 | mouse | 1:200 |
| Mucin 5AC | Abcam | 218363 | rabbit | 1:200 |
| CC16 | Hycult Biotech | HM2178 | mouse | 1:50 |
| Acetylated α-Tubulin | Sigma Aldrich | T6793 | mouse | 1:200 |
| CD144 | Thermo Fisher Scientific | 14-1449-82 | mouse | 1:200 |
| Vangl1 | Sigma Aldrich | HPA025235 | rabbit | 1:100 |

**Supplementary Video 1:** 3D imaging from top to bottom of the Airway Lung-Chip with primary differentiated bronchial epithelial cells cultured in the top channel and microvascular endothelial cells in the bottom channel at 14 days after introduction of the air-liquid interface. Endothelial cells are stained with phalloidin (red) and the epithelial cells with ZO-1 (green) and nuclei with dapi (blue).

**Supplementary Video 2:** 3D imaging of the Airway Lung-Chip with focus on the bottom channel compartment with primary differentiated bronchial epithelial cells cultured in the top channel and microvascular endothelial cells in the bottom channel at 14 days after introduction of the air-liquid interface. Endothelial cells are stained with phalloidin (red) and the epithelial cells with ZO-1 (green) and nuclei with dapi (blue).

**Supplementary Video 3:** Ciliary beat was recorded via phase contrast videomicroscopy. Comparison between chip and insert cultures of the same donor shows no obvious difference at day 14 ALI.

**Supplementary Video 4:** Mucus clearance was visualized by the motion of suspended fluorescent microparticles (here shown as accumulative timelapse to show the trajectories). Chip cultures show more pronounced clearance than inserts of the same donor at day 14 ALI, but at d35 ALI the inserts exhibit well developed clearance. This shows that mucociliary maturation is accelerated on chip compared to insert cultures.
